# Identification of differentially expressed novel tau associated proteins in transgenic AD model of *Drosophila* - a proteomic and *in silico* Approach

**DOI:** 10.1101/2022.05.16.492076

**Authors:** Brijesh Singh Chauhan, Somenath Garai, Jyotsna Singh, Saripella Srikrishna

**Author notes:** Corresponding Author: Saripella Srikrishna^1*^, Cell and Neurobiology Laboratory, Department of Biochemistry, Institute of Science, Banaras Hindu University, Varanasi-221005, Uttar Pradesh, India; Co-corresponding Author: Somenath Garai^2*^, ^2^Department of Chemistry, Institute of Science, Banaras Hindu University, Varanasi-221005 Uttar Pradesh, India.

## Abstract

Alzheimer’s disease (AD) is a memory related neurodegenerative disorder mainly associated with older adults. In this study, transgenic *Drosophila* AD model has been employed to investigate the tau associated proteome. Tau expression was specifically induced in the eye tissues and diseased fly heads were considered for proteomic studies with appropriate controls. We have identified 6 novel proteins from tau induced AD group by 2D and PD Quest analyses and further characterized them by *in-vivo* and *in-silico* approaches. The novel Tau interactors, [Obp44a Isoform A, Pglym Isoform A, IP15846p (Adh variant), RE45450p (mRpL2), Retinin, and Glob1 Isoform B], identified by MALDI-TOF/MS were validated through q-RT-PCR. The altered metabolic, behavioral and mitochondrial dynamics associated with Tau over expressing AD flies could be due to the altered expression of Odorant binding protein 44a (Obp44a) and mitochondrial ribosomal protein L2 (mRpL2), respectively. Further, we have showed for the first time the newly identified protein interaction with tau and other regulatory proteins through protein-protein docking, biocomputational classification and evolutionary relationship using *in silico* studies. Moreover, the study highlights the plausible role of these novel proteins in pathophysiology of phospho-tau induced AD flies and further help unraveling molecular pathways implicated in tauopathies.

## INTRODUCTION

AD, essentially a memory disorder, ensues progressive loss of the cognitive functions in the aged population (Selkoe, 2001; Chauhan et al., 2020). The disease is characterized by extracellular Amyloid-β peptide depositions as senile plaques (SPs) and hyperphosphorylated tau in the form of intracellular neurofibrillary tangles (NFTs). These NFTs are composed of filaments and the main constituent of these filaments is the microtubule associated protein (MAP), Tau. Six isoforms of tau exist through alternative splicing of transcripts from the *MAPT* gene. Splicing of exons 2, 3 leads to 2N at N-terminal and inclusion of exon 10 leads to 4R at C-terminal regions, which forms the longest tau isoform, 2N4R (441 amino acids) in the human central nervous system (Götz et al., 2019; Silva and Haggarty, 2020). From the pathomechanistic point of view, Tau protein is distributed throughout the neuronal cells in culture and upon maturation, it is enriched in the axonal regions of neurons (Kosik and Finch, 1987; Götz et al., 2019).

In healthy cells, tau protein is involved in polymerization and stabilization of microtubules, associated with axonal transport of sub-cellular organelles (Gendron and Petrucelli, 2009; Dolan and Johnson, 2010). The phosphorylation of tau protein declines its binding affinity with tubulin subunit of microtubules and also enhances the self aggregation and fibrillization of phospho-Tau proteins (Cohen et al., 2011; Cisek et al., 2014; Singh et al., 2015), which in turn affects the axonal transport of mitochondria (Ittner and GLJtz, 2011; Mondragón-Rodriguez et al., 2013; Mietelska-Porowska et al., 2014). The disease is caused due to defects in axonal transport or unknown etiological factors, which are yet to be explored. Not much is explored in Tau specific proteome till date. Hence, in order to identify the novel protein candidates involved in disease progression, transgenic *Drosophila* AD model has been employed to study the tauopathies specific proteome. To generate the *UAS-tau-lacZ/ey-GAL4* flies, a cross was made to achieve overexpression of tau between a *ey-GAL4/CyO* driver line and responder *UAS-tau-lacZ* line. The heads of *UAS-tau-lacZ/ey-GAL4* transgenic flies and control *OregonR^+^* flies were taken for proteomic studies. Six differentially expressed proteins, Obp44a Isoform A, Pglym Isoform A, IP15846p (Adh variant), mRpL2, Retinin, and Glob1 Isoform B, were identified by MALDI-TOF. Among them, mitochondrial ribosomal proteins (MRPs) and Odorant binding protein (Obps) exhibited less studies regarding AD pathogenesis (Stamps et al., 2013; Lunnon et al., 2017; Yoo et al., 2017). Healthy mitochondria play a key role in neuronal cell survival through regulation of energy metabolism. Mitochondrial dysfunction and oxidative damage play a major role in AD pathogenesis (Moreira et al., 2010). Several studies reported that AD pathogenesis could be the major consequence of mitochondrial dysfunction, oxidative stress, weight loss and dietary dysfunction (Sultana et al., 2007; Smith and Greenwood, 2008; Moreira et al., 2010; Kai et al., 2015). Mitochondrial dysfunction was found in invertebrate engaged with a relatively short lifespan and progression of cognitive impairment during AD pathogenesis (Bishop et al., 2010; Swerdlow, 2011; Picard and McEwen, 2014). Hence, a detailed study of mitochondrial ribosomal protein L (mRpL2) and Odorant binding protein 44a (Obp44a) was taken up to investigate if they have a role to play in the development and progression of AD.

The normal function of mRpL2 in mitochondrial translation activities, accounts for about 75% constituents of Ribosomal components. Previous reports suggested that deterioration of mitochondrial translation affects the oxidative phosphorylation (OXPHOS) (Fernandez-Ayala et al., 2010; Boczonadi and Horvath, 2014; Brown et al., 2014). The tau pathogenesis could be the consequence of reduced expression of mRpL2 protein leading to mitochondrial translation defect in AD model of *Drosophila* although exact mechanism is not known. The alteration in mitochondrial morphology and membrane potential is characteristic feature of mitochondrial damage associated with failure of multiple respiratory chain machinery resulting in decreased adenosine triphosphate (ATP) levels in the brain tissues (Cai, 2017).

Therefore, detailed study associated with mitochondrial dynamics is required for understanding the expression of mRpL2 and its association with the mitochondrial fission and fusion events in Tau induced AD tissues. The mitochondrial markers such as Dynamin-related protein-1 (Drp1) is involved in regulation of mitochondrial fission, Mitochondrial assembly regulatory factor-1 (Marf1) plays role in mitochondrial fusion, Parkin (ubiquitin ligase) is required for degradation of damaged mitochondria and HtrA1/2 (mitochondrial serine protease) involved in caspase-dependent apoptosis. All these markers regulate the fission and fusion process of mitochondrial membrane (Yadav and Srikrishna, 2019; Chauhan et al., 2021). A recent study suggested that hyperphosphorylated tau protein is involved in mitochondrial dysfunction during AD progression. The hyperphosphorylation or abnormal tau protein aggregation impedes mitochondrial distribution due to failure of Axonal transport (Cheng and Bai, 2018). In this context, the newly identified mRpL2 and Obp proteins gain significance in relation to the dysfunction of mitochondria in Tau induced AD pathogenesis. Therefore, reduced expression of mRpL2 during AD progression has emerged as new findings in AD model of *Drosophila.* Previous studies showed that the mutation/alteration in mitochondrial ribosomal proteins (MRPs) decreases the life span of *C.elegans* due to increasing sensitivity to oxygen (Fujii et al., 2011; Mozhui et al., 2017). Therefore, longevity assay is also needed to ascertain the life span of tau induced AD flies. Similar study on the life span of tau based *Drosophila* models of AD was also carried out in line of the present studies (Wittmann et al., 2001; Colodner and Feany, 2010; Higham et al., 2019).

Odorant binding proteins (Obps) are water soluble low molecular weight proteins abundantly present in the lymph of chemosensory sensilla. They are essential for transporting hydrophobic odorants through olfactory sensilla containing aqueous lymph to olfactory receptor neurons, which subsequently convey signals to the *Drosophila* brain in response to the odor. The olfactory sensilla contain olfactory receptor neurons (ORNs) submerged in the aqueous lymph. The Obp present in the olfactory sensilla, a taste organ in the fruit flies, is capable of perceiving the food odor (Larter et al., 2016). Recent information suggests that alteration in expression of olfactory proteins is involved in the development of AD pathogenesis (Yoo et al., 2017; Archunan, 2018). In *Drosophila* there are approximately 51 odorant binding proteins present comparable to its odorant-receptors (Hekmat-Scafe et al., 2002). In present study the down regulation of novel Obp44a protein observed in tau induced AD model could be responsible for increased anorexia symptoms, which reflected in reduced food intake and loss of body weight in Tau induced AD flies. Interestingly, the present study sheds some light on the functional relationship of mRpL2 and Obp44a proteins with tau, although there is no clear evidence to suggest the direct relationship of mRpL2 and Obp44a with tau protein otherwise. Further, the detailed study on these newly implicated proteins in relation to Tau would help in unraveling yet unidentified and underlying mechanisms of Tau mediated AD pathogenesis.

## MATERIALS AND METHODS

### Rearing and Maintenance of Fly Stocks

The Fly Stocks *OregonR*^+^, *ey-GAL4/Cyo, UAS-GFP,* and *elavc155-GAL4* lines were obtained from the Bloomington *Drosophila* Stock Center, Indiana, USA and fly stock *UAS-tau-lacZ,* a kind gift by Dr. Andrea Brand, University of Cambridge, United Kingdom. The flies stocks, *OregonR*^+^, *ey-GAL4/CyO and elavc155-GAL4* served as control and *UAS-tau-lacZ* driven with different GAL4 driver lines served as AD disease model in all experiments. Flies were cultured in standard corn meal agar media in BOD incubator at 24±1°C on a 12 hr light/ dark cycle, where specific conditions applied which can enhance the GAL4 activity in flies raised at 28±1°C.

### X-Gal Staining

The eye imaginal discs of 3^rd^ instar larvae of *UAS-tau-lacZ/ey-GAL4* flies were dissected in a freshly prepared Poels’ salt solution (NaCl 86 mg, KCl 313 mg, CaCl_2_.2H_2_O 116 mg, NaH_2_PO_4_.2H_2_O 88 mg, KHCO_3_ 18 mg, MgSO_4_.7H_2_O 513 mg, ddH_2_O upto 100 ml and pH was adjusted to 7.0 with 1M NaOH). Tissues were washed with 50 mM Sodium-phosphate buffer (wash buffer, pH 8.0) for 3 times. Subsequently, tissues were fixed in 2.5% Glutaraldehyde for 10 min at room temperature (RT). The fixative was removed and the tissues were washed three times with wash buffer (50 mM Sodium-phosphate) for 10 min each. Afterward, 400 μl X-gal staining solution (60μl of 5% X-gal in Dimethyl formamide, 20μl of 100mM K_3_ [Fe (CN) 6], 20μl of 100mM K_4_ [Fe (CN) 6], 50μl of 1M Sodium phosphate buffer, and 850μl dd H_2_O) was added and incubated at 37°C in dark for 40 min to develop the blue color. X-gal stain was removed and tissues were washed three times with wash buffer for 10 min each. The tissues were then transferred on to a fresh slide and mounted with 80% glycerol, followed by observation under bright field microscope.

### Specimen Preparation for Scanning Electron Microscopy (SEM)

The 15 day old, age matched, *OregonR^+^* and tau induced AD flies were taken for SEM studies. The sample preparation with minor modifications was done as described by (Wolff, 2011). The flies were immersed in 1ml fixative [(1% Glutaraldehyde (25μl), 1% formaldehyde (0.4ml), 0.1M phosphate buffer pH 7.2 (100μl) and ddH_2_O (0.475 ml)] for 2h at room temperature (RT). One small drop of 0.2% tritonX-100 was also added to reduce the surface tension, so that flies can easily be submerged in the fixative. After incubation, flies were rinsed with ddH_2_O and dehydrated by adding a series of ethanol treatment in gradient manner (once in 25%, 50%, 75% and twice in 100% ethanol) at RT for 12h. Afterwards, flies were coated with sputter-coat gold palladium alloy to produce high topographic contrast and resolution of fly eye and olfactory organs using ZEISS EVO 18 SEM.

### Proteomics Study

The head tissues of *OregonR^+^* and *UAS-tau-lacZ/ey-GAL4* transgenic lines were processed as under for proteomic studies to identify the differentially expressed novel candidate proteins.

### Extraction of Protein

In order to isolate total proteins, heads of wild type and experimental (*UAS-tau-lacZ/ey-GAL4*) flies (60 each) were homogenized in lysis buffer: 20 mM Tris-HCl (pH 7.6) and 1μl/ml PMSF cocktail. The homogenized samples were centrifuged at 10,000 rpm at 4°C for 20 min and supernatants were collected in fresh eppendorf tubes. The protein concentration was quantified by Bradford method (Bradford, 1976). The protein lysate was stored at -20°C until further use.

### Procedure for 2D GEL Electrophoresis

Equal amount of protein concentrations (150μg) were taken to rehydrate the BIO-RAD IPG pH gradient strip (pH range 3-10) for 16-17 hr. Rehydrated BIO-RAD IPG strips were used for Iso- Electric Focusing (IEF) in BIO-RAD IEF machine for 23 hr. After completion of IEF program, the strips were treated with equilibrium buffer: 6M Urea, Tris-buffer (0.375M), 2% SDS, 20% Glycerol for 15 min each. Equilibrium buffer [1] had 0.2% DTT and equilibrium buffer [2] had 2.5% Iodo-acetamide. After that the strips were processed for gel electrophoresis in 12% resolving gel at 100 volt.

After completion of gel run, the gels were fixed in fixative (Methanol 150 ml, Acetic acid 36 ml, Formalin 150 μl, ddH_2_O 114 ml) on shaker for overnight. Gels were washed thrice with 20% ethanol for 20 min each, at the end of which sensitizing solution (double distilled water (ddH_2_O) 300 ml, Na_2_S_2_O_3_.5H_2_O 0.06 g) was added for 2 min. The gels were washed twice with ddH_2_O for 1 min. Subsequently, prechilled AgNO_3_ solution (ddH_2_O 300 ml, AgNO_3_ 0.6 g and HCHO 228 μl) was added for 30 sec. After this treatment, AgNO_3_ solution was removed and developing solution (6% Na_2_CO_3_, 0.0004% Na_2_S_2_O_3_.5H_2_O, and 0.05% HCHO) was added to develop the spots on gels via gentle shaking. Again gels were washed twice with ddH_2_O and the development was terminated by adding termination solution. Images of gels were captured by molecular gel doc (BIO-RAD GEL/CHEMI DOC, Japan).

### Analysis of Protein Spots on 2D GEL Image

PDQuest software (PDQuest basic 8.0.1, BIO-RAD) was used to determine the difference in protein expression profile from spot developed gels. Three replicate images of Control and test were used for automatic spot detection using PDQuest 2D gel analysis software. The spot intensities of these replicates from control and test were subjected to independent analysis by Standard spot number (SSP). The protein spots showing the altered expression in spot quantity between control and test groups were marked in 2D-gels and significant spots were excised for MALDI-TOF/MS analysis.

### Trypsin Digestion of Protein Spots

The gel slices were diced to small pieces and placed in eppendorf tubes and were destained using destaining solution (1:1 Ratio of ACN and 25mM Ammonium Bicarbonate) with 10 minute intervals (3-4 times) until the gel pieces become translucent white. The gels were then dehydrated using Acetonitrile (100µl) and thermo-mixed at 600 RPM for 10 minutes till complete dryness. Then they were rehydrated with 10mM DTT (100µl) and incubated for 60 min. After incubation, the DTT solution was removed. The gel pieces were now incubated with 55mM Iodo-acetamide (100µl) for 45min. The supernatant was discarded and the gel was incubated with ammonium bicarbonate solution (100µl) for 10min. After that, supernatant was removed and the gel was dehydrated with Acetonitrile (100µl) for thermo-mixed at 600RPM for 10 min till gel was de-hydrated. Then Trypsin solution (8µl) along with 150µl of 25mM Ammonium Bicarbonate was added and incubated overnight at 37°C. The digested solution was transferred to fresh eppendorf tubes. The gel pieces were extracted thrice with extraction buffer (1:1 ratio of ACN and 0.1% Triflouroacetic acid) and the supernatant was collected each time into an eppendorf and then Speedvacc’d till complete dryness. The dried pepmix was suspended in TA buffer (0.1% TFA and Acetonitrile in the ratio of 2:1) prior to MALDI analysis.

### Matrix Assisted Laser Desorption Ionization-Time of Flight /Mass Spectrometry (MALDI-TOF/MS)

The peptides obtained were mixed with HCCA (α-Cyano-4-hydroxycinnamic acid) matrix (5 mg/mL α-Cyano-4-hydroxycinnamic acid in 1:2 ratio of 0.1% TFA and 50% ACN) in 1:1 ratio and the resulting 2μl was spotted onto the MALDI plate [(MTP 384 ground steel (Bruker Daltonics, Germany)]. After air drying the sample, it was analyzed on the MALDI TOF/TOF ULTRAFLEX III instrument (Bruker Daltonics, Germany) having Nd : YAG smart laser beam and external calibration was done with standard peptide (PEPMIX Mixture) supplied by Bruker, with masses ranging from 16107 to 509304 Da. Further analysis was done with FLEX ANALYSIS SOFTWARE (Version 3.3) in reflectron ion mode with an average of 500 laser shots at mass detection range between 500 to 5000 m/z for obtaining the PEPTIDE MASS FINGERPRINT. The masses obtained in the peptide mass fingerprint were submitted for Mascot search in “CONCERNED” database for identification of the protein. Protein scores were derived from ions scores as a non-probabilistic basis for ranking protein hits. We had found maximum 20 hits and uses the top protein scores greater than 70 were exhibited significant (*p<0.05*) (Table 1).

**Table 1:**
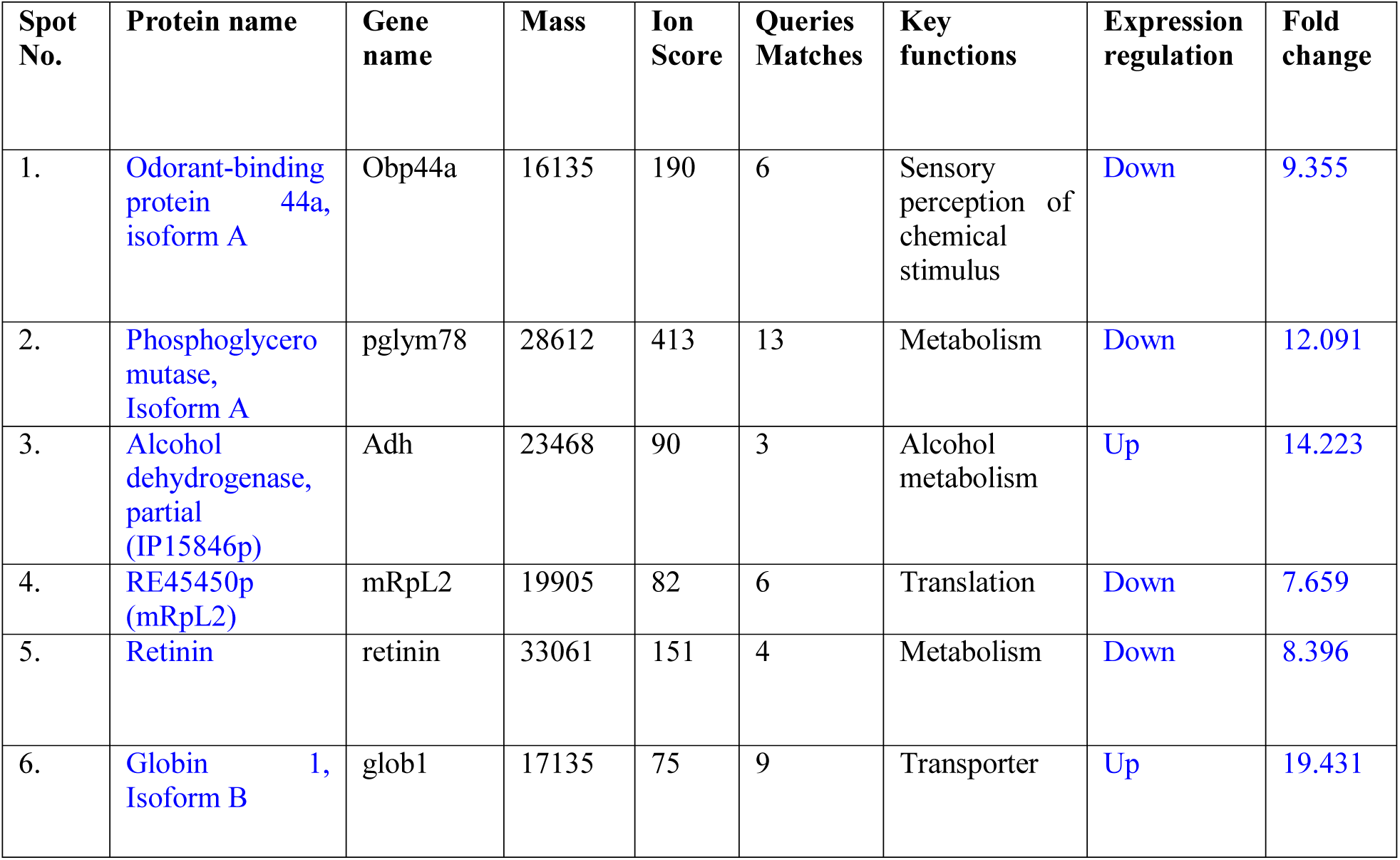
Differentially regulated six novel proteins from tau induced AD model of Drosophila.

### RNA Isolation, cDNA Synthesis and Quantitative Real Time PCR (qRT-PCR) and Heatmap Analysis

The mRNA level of Obp44a Isoform A, Pglym Isoform A, IP15846p (Adh variant), mRpL2, Retinin, and Glob1 Isoform B were analyzed using qRT-PCR. For RNA extraction 70 fly heads each were harvested from *OregonR^+^* and *UAS-tau-lacZ/ey-GAL4* flies. Here after, followed the prescribed RNA isolation method mentioned by (Yadav and Srikrishna, 2019). High quality RNA concentrations were measured using ultraviolet spectrophotometer at OD260/280 ratio. Further, cDNA were synthesized using (1000nM) extracted RNA through Reverse transcriptase PCR (Sambrook and MacCallum, 2001). These cDNA from both samples were used as template DNA for quantitative expression of Obp44a Isoform A, Pglym Isoform A, IP15846p (Adh variant), mRpL2, Retinin, and Glob1 Isoform B normalized against gapdh. Further, relative mRNA expression of mitochondrial dynamics regulators were performed using cDNA of the respective samples as template DNA for quantification of *Drp1, Marf1, HtrA1/2, and parkin* normalized against *Gapdh.* In addition, relative mRNA expression of *Obp99b* and tumor suppressor gene *p53* was also performed using cDNA of respective samples as template DNA normalized against gapdh using following set of primers (Table 2).

**Table 2:**
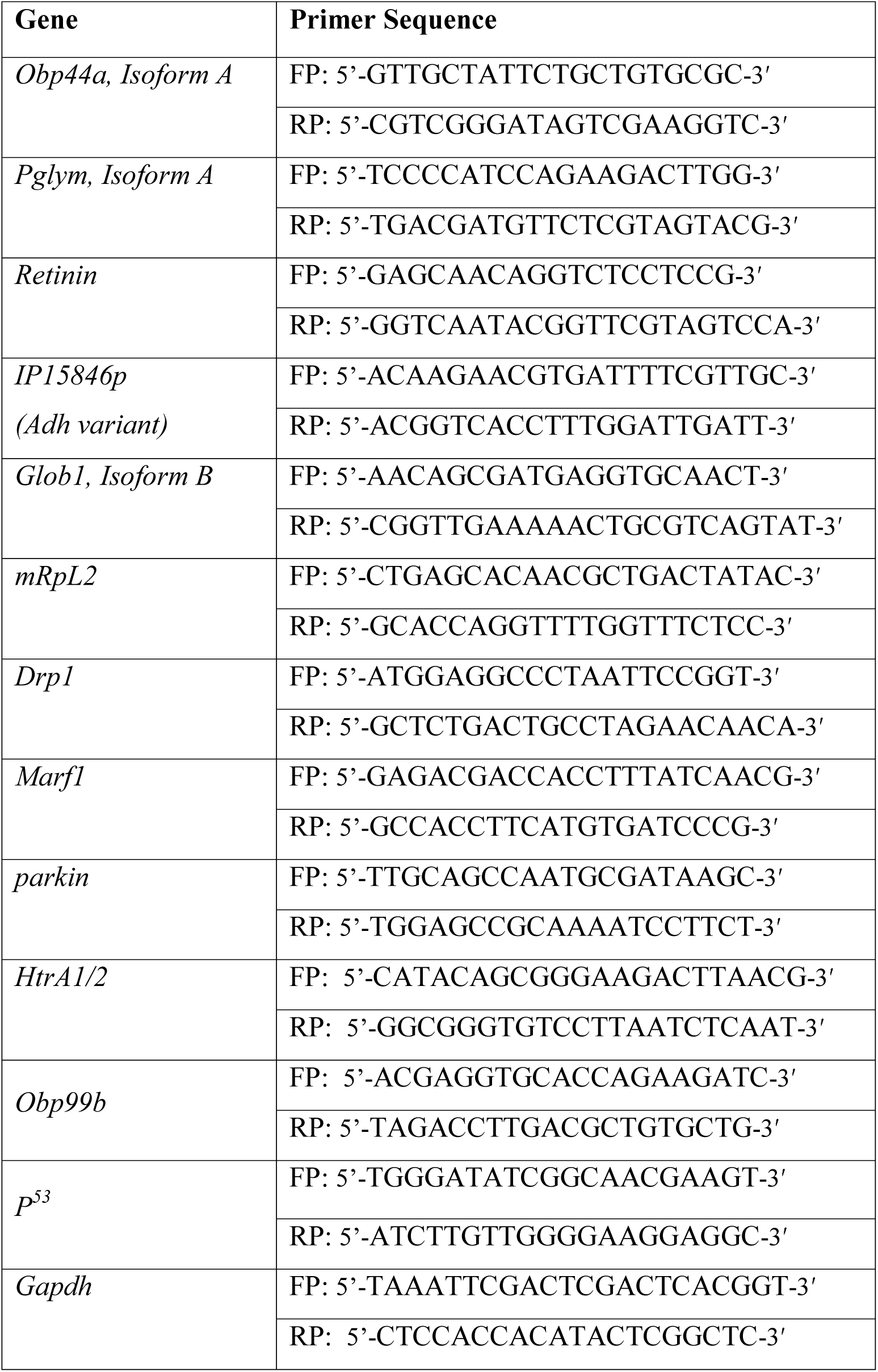
Primers for newly identified genes were used for qRT-PCR validation experiments and also for study of mitochondrial dynamic regulators.

The intercalating dye SYBR green JUMPSTART TAQ Ready mix (Thermo Fisher Scientific) used for quantitate the mRNA expression level using qReal-Time PCR (Applied Biosystems 7500). All samples were performed in triplicates and relative fold changes were calculated using comparative ΔΔCт values. Further, expression heatmap of novel identified genes were built up using freely available web server http:/biit.cs.ut.ee/clustvis/. In which unit variance scaling is applied to rows; Singular value decomposition (SVD) with imputation is used to calculate principal components.

### Analysis of Mitochondrial Morphology and Membrane Potential

To examine the mitochondrial morphology, *ey-GAL4>UAS-mitoGFP* as control and *ey- GAL4>UAS-tau-lacZ* + *UAS-mitoGFP* as test larval lines have been used to express GFP tagged mitochondria endogenously. The 3^rd^ instar larvae of these lines were dissected in 1xPBS (pH 7.4) to isolate eye imaginal discs, followed by incubation with 300nM Mito-Tracker-Red (Thermo Fisher-Scientific) at RT in dark for 45 min. Thereafter, tissues were washed quickly twice with 1xPBS and fixed with 4% paraformaldehyde at RT for 15 min. Afterward, tissues were washed 2 times with 1xPBS for 5 min each and subsequently mounted on glass slides using DABCO. The images were captured by the confocal microscope (Zeiss LSM-510 Meta) and fragmentation and elongation of mitochondria analyzed manually.

### ATP Quantitation Assay

The 15 days old fly heads of wild type (*OregonR^+^)* and *UAS-tau-lacZ/ey-GAL4* were decapitated with the aid of fresh needle. The fly heads weighing 10 mg were taken from control and disease group separately in 1.5 ml eppendorf centrifuge tube for ATP quantitation. The content of ATP was measured using ATP colorimetric assay kit (Bio-Vision, Catalog # K354-100, USA) as per manufacturer’s guidelines. Further tissues were homogenized using homogenizer on ice and subjected to deproteinization of sample adding 25μl Trichloroacetic acid (Sigma Aldrich, Catalog # T6399). Finally, the ATP content was normalized against the protein concentration using Bradford method (Bradford, 1976). The experiment was performed in triplicate and data was analyzed using student’s t test.

### Food Intake, a Y-maze Assay and Body Weight Measurement Analysis

Food intake and body weight measurement analysis was performed in age matched *OregonR^+^* and *UAS-tau-lacZ/ey-GAL4* flies. To perform the food intake tracer experiment, 15 days old flies from both groups were initially subjected to fasting for 1hr in BOD incubator at 28±1°C and followed the food intake procedure with minor modifications as described by (Sun et al., 2013) . Afterward, 5 flies from both groups were transferred separately in cornmeal food vials containing 0.5% Trishul green synthetic dye. At the same time, similar numbers of control flies were also transferred in vials containing normal food media. The experiments were setup in triplicate and flies were allowed to feed for 30 min. After that, their heads were decapitated from body part with fresh fine needle and homogenized the fly bodies in 200μl distilled water with plastic pestle and followed by addition of 800μl distilled H_2_O. The homogenate was centrifuged at 12,000 x g for 2 min and 0.9 ml supernatant transferred in fresh new eppendorf tube. After that, volume was maintained 1.5 ml by adding distilled H_2_O and centrifuged again for 2 min. Sample absorbance measured at 630 nm using synergy H1 Multimode plate reader, Biotek. The control flies absorbance used to correct for background absorbance of flies and food intake calculated using net absorbance. Y-maze assay was also performed for testing the olfactory response of *OregonR^+^* and AD flies (N=150 flies in each group) (Simonnet et al., 2014). The experiments were performed in triplicates. Olfactory index in percentage was calculated using the following formula.

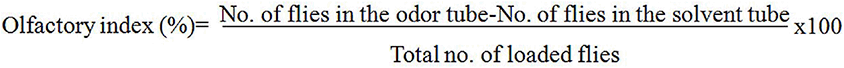

Further, weight measurement experiment was performed using age matched male and female flies from *OregonR^+^* and *UAS-tau-lacZ/ey-GAL4*, respectively. The 15 day old tau induced flies (50 male + 50 female) were taken for weight measurement. Similarly, same method also applied for *OregonR^+^*flies. All experiments were performed in triplicates and data was analyzed using student’s t-test.

### Cell Viability Assay

To examine the metabolic activity of *Drosophila* eye imaginal discs, primerly *UAS-tau-lacZ/ey- GAL4-UAS-GFP* and *OregonR^+^*3^rd^ instar larvae were chosen for experiment. The eye imaginal discs from both groups (10 pairs in each group) were dissected in 1xPBS (pH 7.4) separately in triplicate for viability assay. The tissues were washed twice with 1xPBS and incubated in 100μl of 0.6 mg/ml MTT [3- (4, 5-dimethylthiazol-2yl)-2, 5-diphenyl tetrazolium bromide] (MP Biomedical, France) for 2 hr at 37 °C. After end of incubation, MTT solution was removed and tissues were washed twice with 1xPBS. Afterward, 200 μl DMSO (Sigma Aldrich) was added for 1 hr at 37 °C to solubilize the insoluble formazan crystals. The colored solutions were transferred into 96 well culture plates. The color intensity was measured using multimode plate reader (Synergy H1 Biotek) at 570 nm wavelength and data was analyzed using student’s t-test.

### Life Span Measurement

The survival assay was performed to measure the life span of fruit flies. For experiment 10 flies were taken in each vial. There were three groups’ elavc155-GAL4, OregonR*^+^* and human tau protein expressing flies cultured in standard cornmeal media at 28±1°C in BOD incubator. Three vials have been taken for each group (n=30). The *OregonR^+^* and *elavc155-GAL4* considered as control while *elavc155-GAL4>UAS-tau-lacZ* flies as test. The reckonings of viable flies were done every day. Assays were conducted in triplicates. The differences in survival data analyzed using the Log-rank (Mantel-Cox) Test.

### Protein-Protein Docking Methodologies

The selected protein sequences of the identified proteins from MALDI-TOF/MS were downloaded from Alpha Fold Protein Structural Database (BIOVIA, Dassault Systèmes, 2020; Jumper et al., 2021) for the organism *Drosophila melanogaster.* The Structures were optimized for protein-protein docking studies with the help of Biovia Discovery Studio Visualizer from the Dassault systems (BIOVIA, Dassault Systèmes, 2020). The finalized PDB files were subjected for protein-protein docking studies through the GRAMM-X Protein Docking Web Server (Tovchigrechko and Vakser, 2005; Tovchigrechko and Vakser, 2006). The combined complex networks of protein-protein docked structures with multiple orientations of the coupled proteins were thereafter analyzed with PDBsum Generate structural analysis server from EMBL-EBI (Laskowski et al., 2018). The complete protein-protein interactions like hydrogen bonding schemes, salt bridge interactions, non-bonding interactions, identification of the schematic interaction motifs and detailed Ramachandran plots with the corresponding types/numbers were all generated by the PDBsum analysis. The figures for the protein-protein interactions were developed by subsequent use of the Biovia Discovery Studio Visualizer and PyMOL visualization system by Schrödinger under the academic research license from Banaras Hindu University (Schrödinger and DeLano, 2020). The post-analysis figure management was performed through available online resources. The complete protein-protein docking results were represented in the Figure 9 and Figure 10, Tables S1, S2, and S3 accordingly.

### *In Silico* Analysis of Protein-Protein Interaction and Functional Classification

The network analysis of individual protein obtained from MALDI-TOF was depicted through flybase interaction browser (www.flybase.org/cgi-bin/get_interactions.pl). The interaction browser provided additional essential information about the complex interactions of individual protein with other proteins in the head region of AD flies. Further, network interaction of individual identified protein was build via online esyN network builder software (www.esyn.org/). Further, molecular function, biological process, and protein class of identified proteins from MALDI-TOF/MS were analyzed using online bioinformatics tool Panther Version 13.1 (www.panther.org) (Mi et al., 2013).

### Molecular Phylogenetic Study of Identified Proteins

The evolutionary history was inferred using the UPGMA method (Sneath and Sokal, 1973). The tree is drawn to scale, with branch lengths (next to the branches) in the same units as those of the evolutionary distances used to infer the phylogenetic tree. The evolutionary distances were computed using the Poisson correction method (Zuckerkandl and Pauling, 1965) and are in the units of the number of amino acid substitutions per site. The analysis involved 7 amino acid sequences. All positions containing gaps and missing data were eliminated. There were a total of 112 positions in the final dataset. Evolutionary analyses were conducted in MEGA7 (Kumar et al., 2016).

### Statistical Analysis

The statistical information of two groups, control (*OregonR^+^*) and diseased (*UAS-tau-lacZ/ey- GAL4*), acquired from 2D, qRT-PCR, ATP assay, survival assay, weight measurement and food intake assays, were analyzed using Graph Pad Prism-5 software. Experiments were executed in triplicate and values were presented through two tailed t-test. The two way ANOVA and Log-rank (Mantel-Cox) test depending on experimental groups to determine statistical significance. The *p* value *<0.05* reflected as statistically significant.

## RESULTS

### Microscopic Study of *Drosophila* Eye Phenotypes

The eye tissue specific expression of Tau protein was confirmed through X-gal staining in eye imaginal discs of 3^rd^ in star larvae. The expression of *lacZ* reporter in eye imaginal disc confirms the positive activity of β-galactosidase; that reports the presence of *tau-lacZ* fusion protein at the posterior region in eye imaginal discs (Figure1A, D). Further, eye phenotypes of 15 day old adult flies were examined through both light and scanning electron microscopy studies. In wild type, the eyes exhibited normal phenotype (Figure 1B,C) where as ommatidial disruption and rough eye phenotypes were observed in case of *UAS-tau-lacZ/ey-GAL4,* as a consequence of over expression of tau protein (Figure 1E, F). The ommatidial disruption spanning the area of eye morphology measured with the Image J 1.38x tool as shown in Figure 1G.

**Figure 1:**
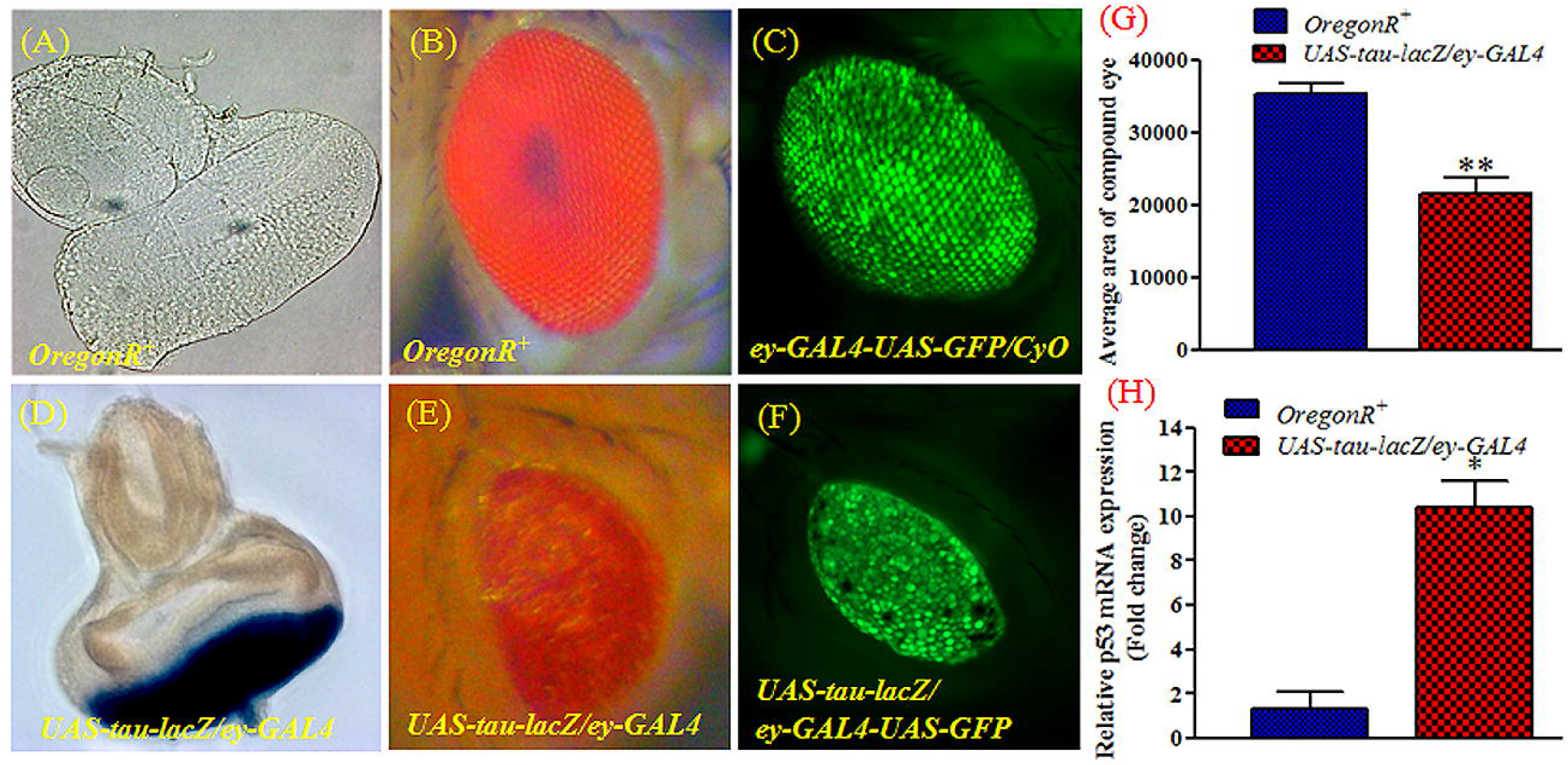
Represents X-gal staining and eye phenotype of Drosophila. (A) X-gal staining of eye imaginal discs of wild type *OregonR^+^* exhibited no lacZ expression, (B) Eye phenotype of wild type *OregonR^+^*, (C) Normal eye phenotype of *ey-GAL4-UAS-GFP/CyO* flies, (D) X-gal staining for eye imaginal discs of *UAS-tau-lacZ/ey-GAL4* shows evidence of Tau protein expression, (E and F) Rough eye phenotypes of *UAS-tau-lacZ* driven by *ey-GAL4* and *ey-GAL4-UAS-GFP* respectively. Histogram (G) represents the reduced eye area (μm^2^) in AD flies in contrast to control, significance ascribed as *\*\*p<0.0018*.

Moreover, scanning electron microscope (SEM) also revealed complete loss of eye bristles and reduced eye size with disturbed patterning of ommatidia as compared to control (Figure 2 I, II). Scanning electron microscopy (SEM) of antennae of OregonR^+^ and tau induced AD flies revealed that the olfactory sensilla of tau flies were poorly developed in comparison with control (Figure 2III).

**Figure 2:**
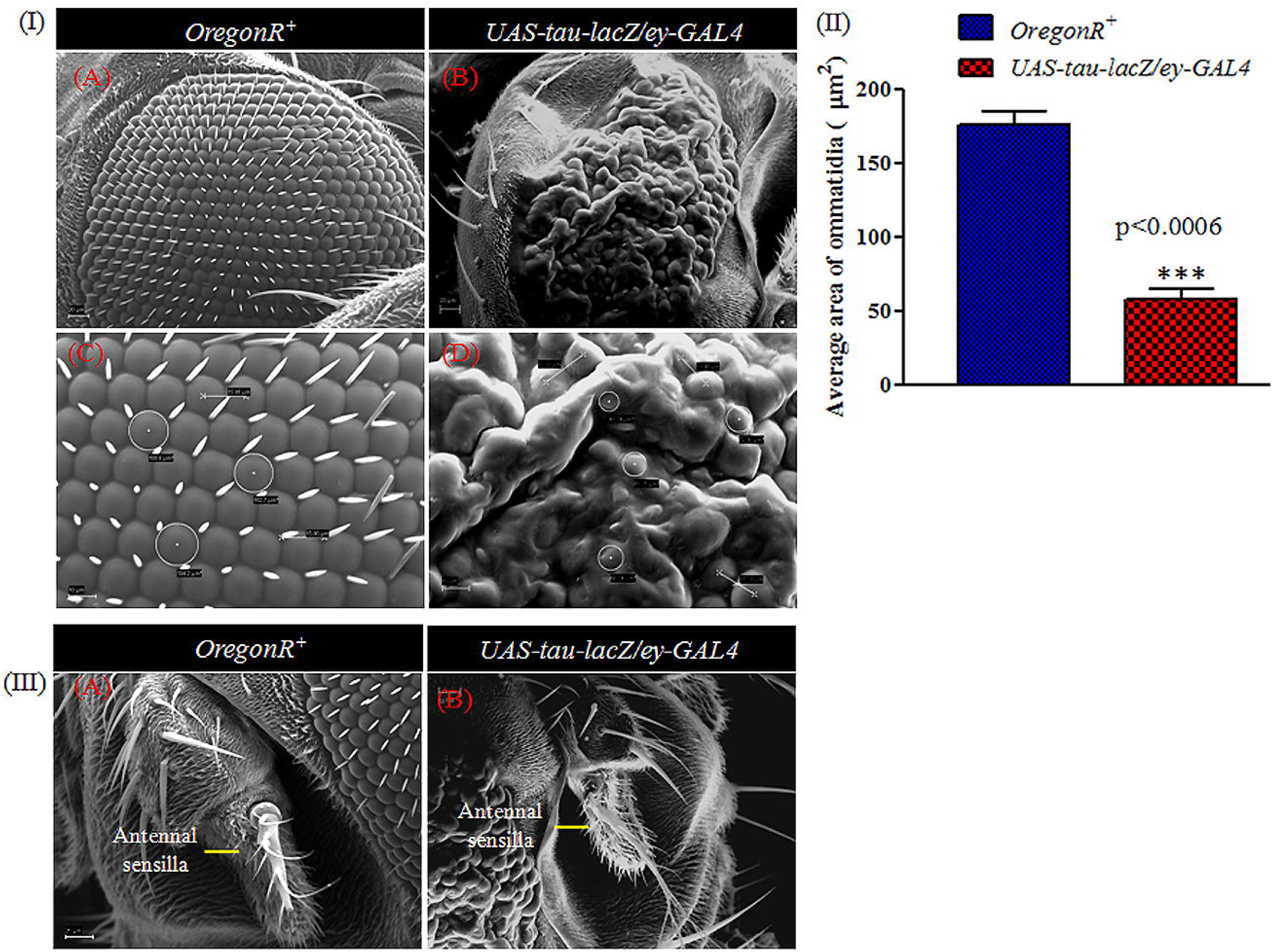
Scanning electron microscope (SEM) shows the compound eye morphology and olfactory organ of *OregonR^+^* and tau induced AD flies. (I) Illustrates the reduced eye morphology, disturb pattern of ommatidia and complete loss of bristles in AD flies as compared to *OregonR^+^*. (II) Histogram represents the reduced area of ommatidia in eye of AD flies compared to OregonR^+^. (III) Shows poor development of olfactory organs in AD flies compared to *OregonR^+^*.

### Differential Proteome Analyses of *Drosophila* Head Proteins

The over expressed tau protein induced changes at proteome level in comparison to wild type was revealed by MALDI-TOF/MS analysis. Protein samples (150μg) isolated from wild type and experimental fly heads was resolved by 2D gel electrophoresis. The resolved protein spots were automatically detected from gels and matched with the PD Quest software basic 8.0.1, BIO-RAD (Figure 3). The protein quantification of individual spots of control and AD sample was done through spot volume and intensity. There were 358 spots identified after automatic quantitative analysis set between control and AD. Of these, six spots showed significant alterations in AD as compared to control (Figure 4). These spots were excised and used for MALDI-TOF/MS analysis. The fold change of identified proteins from MALDI-TOF/MS analysis is given in Table 1. These proteins are Odorant binding protein 44a Isoform A (OBP44a, Isoform A), Phosphoglyceromutase isoform A (Pglym, Isoform A), Retinin, Alcohol dehydrogenase (Adh), Globin 1 Isoform B (Glob1, Isoform B), and mitochondrial ribosomal protein L2 (mRpL2), which are differentially expressed between control and AD group.

**Figure 3:**
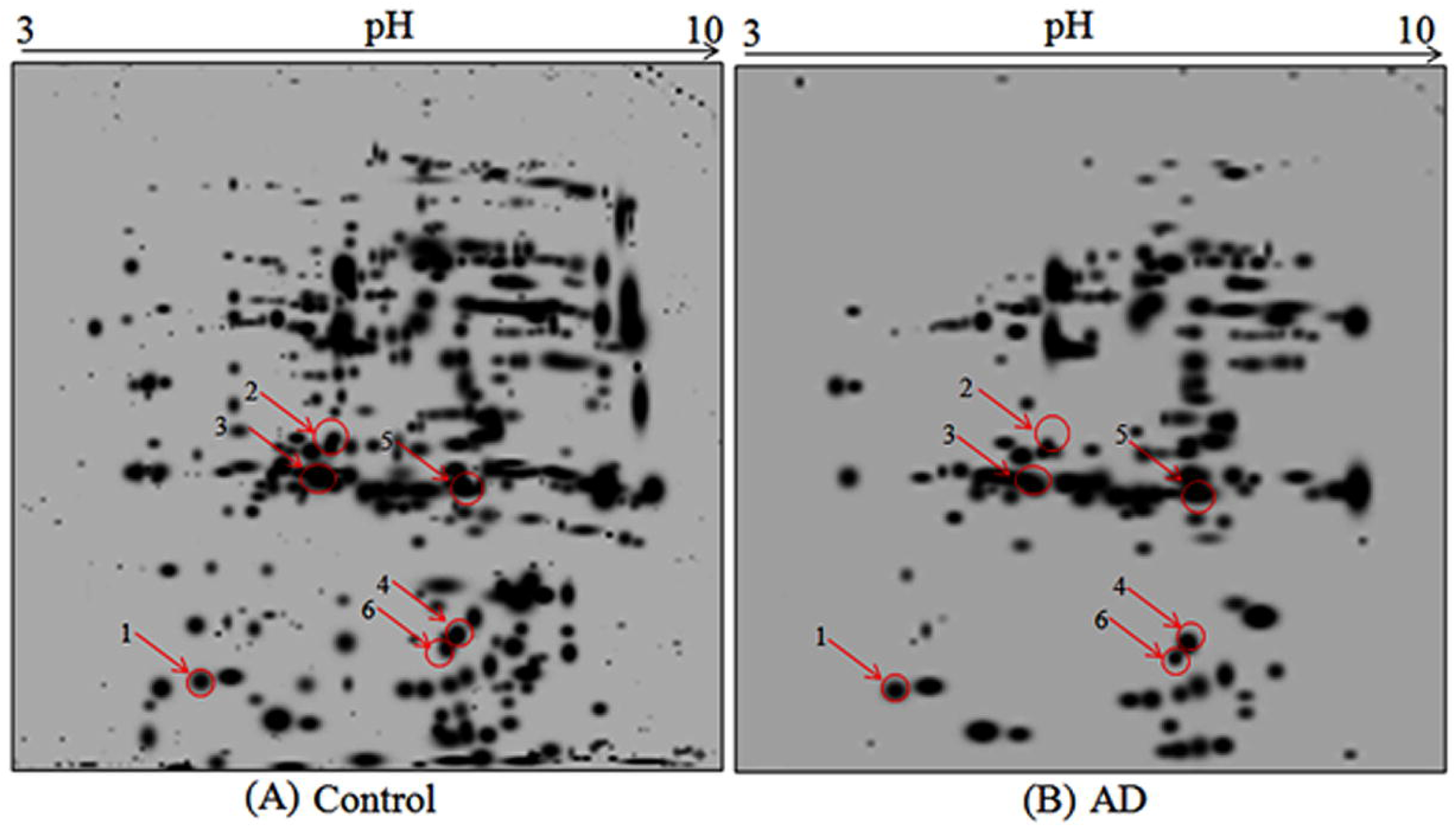
Differential proteome analyses of head proteins of Control *(OregonR^+^)* and AD flies *(UAS-tau- lacZ/ey-GAL4)*. Equal amounts of proteins were separated by 2D gel electrophoresis. (A) 2D gel image of *OregonR^+^*, (B) 2D gel image of transgenic *UAS-tau-lacZ/ey-GAL4*. The pH range of BIO-RAD IPG strip is indicated in the figure. Only differentially expressed protein spots were selected for MALDI-TOF/MS analysis.

**Figure 4:**
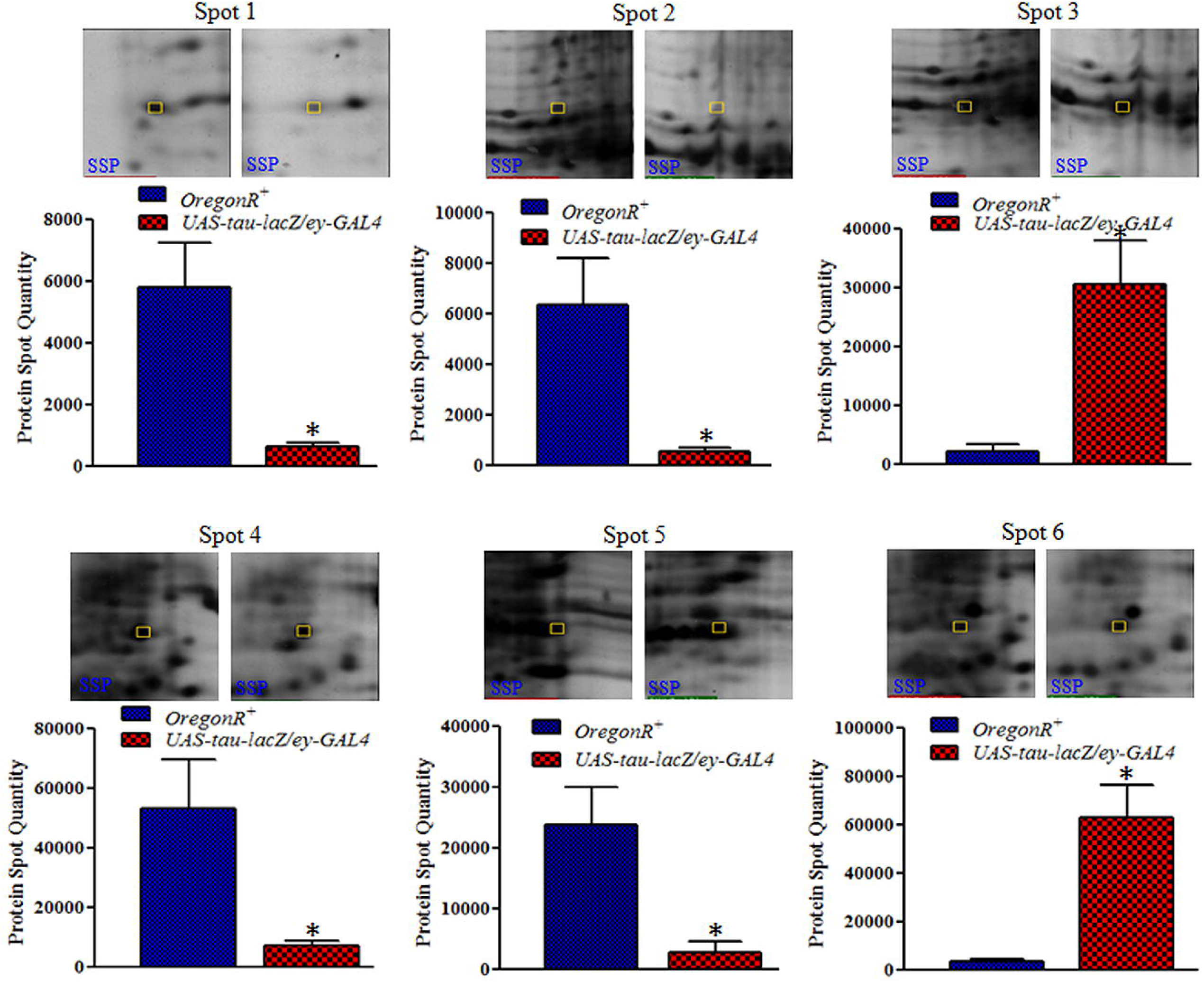
Histogram represents the differentially expressed protein spots shown in magnified views above the graphs (spot 1-6). The protein spot of *OregonR^+^* and *UAS-tau-lacZ/ey-GAL4* were plotted as mean of protein intensity values from three replicates. The significant difference between control and AD is ascribed as *p *< 0.05*. Data is displayed as means + SEM of three paired student’s t-test.

### Validation of Differential Expression of Newly Identified Proteins

The mRNA levels of *tau* associated novel genes identified through MALDI-TOF/MS analysis were assessed through quantitative real time-Polymerase chain reaction (q-RT PCR). The outcome shows that *Alcohol dehydrogenase (Adh)* and *Globin 1(glob1)* exhibit approximately 1.5 and 3.8 fold upregulation of mRNA expression, respectively, while *obp44a*, *pglym78*, *mRpL2* and *retinin* exhibit down-regulation of mRNA expression in AD fly heads, as compared to controls (Figure 5A). Moreover, level of tumor suppressor gene p53 mRNA expression was also measured and was found to have more than 10 fold upregulation in the head tissues of tau flies as compared to wild type, suggesting that overexpression of p53 could be involved in elevation of tau phosphorylation (Figure S1). Similarly, relative mRNA expression of obp99b was also measured and found to be 4 fold down regulated in AD fly heads in contrast to control (Figure S2).

**Figure 5:**
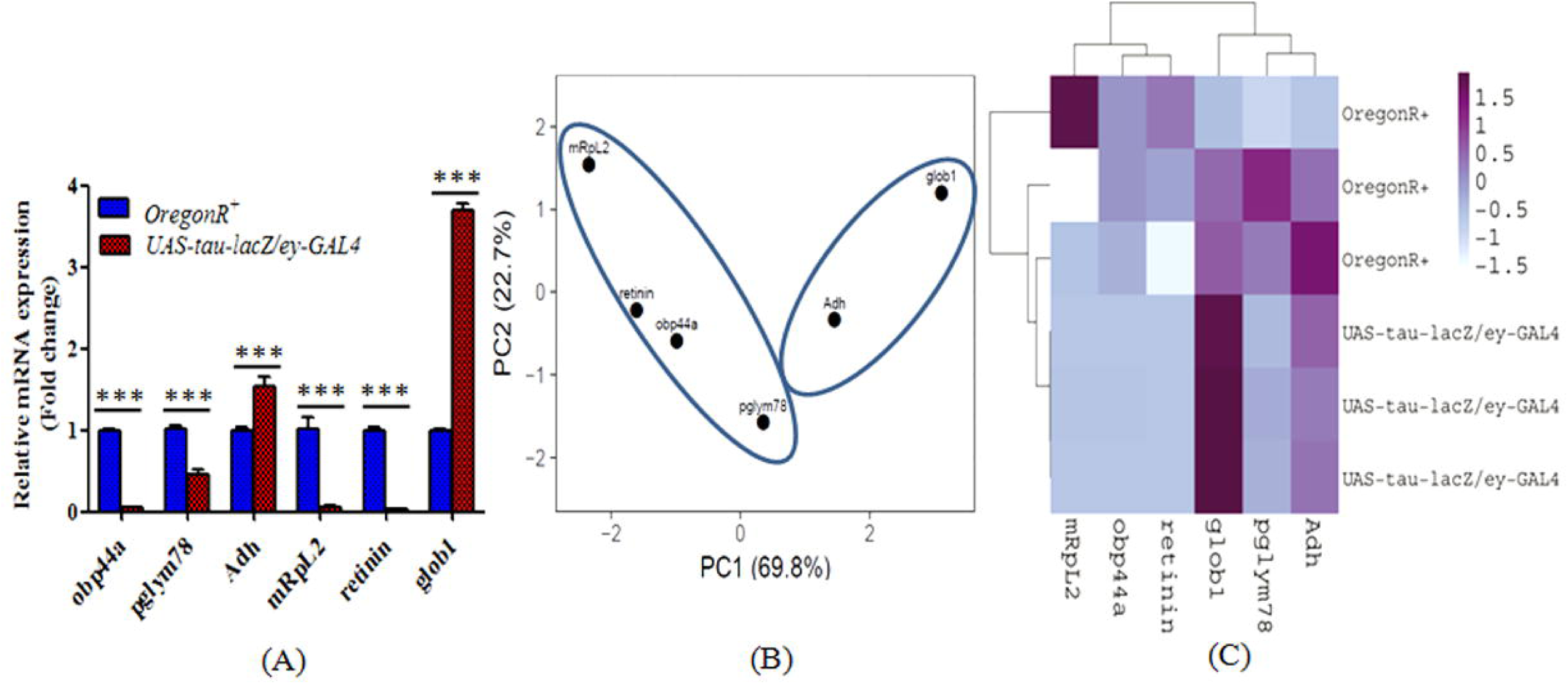
(A) Relative mRNA expressions of novel identified genes in control and experimental samples (B) Principal components, (C) Heatmap of four downregulated [obp44a, Pglym78, mRpL2 and retinin] and two upregulated [IP15846p (Adh) and glob1] genes expressed in the head tissues of *UAS-tau-lacZ/ey- GAL4* as compared to *OregonR^+^* flies. The data were analyzed using 2way ANOVA test *\*\*\*p<0.001*.

Further, expression heat map of respective genes was plotted on X and Y axes which show principal component 1 and principal component 2 of 69.8% and 22.7% of the total variance respectively (N = 6 data points). Further, rows are centered; unit variance scaling is applied to rows. Both rows and columns are clustered using correlation distance and average linkage 6 rows, 6 columns (Figure 5B and 5C, Metsalu and Vilo, 2015). The scattered plot for *Drosophila* tau associated gene expression was visualized in a heatmap. In AD flies, an increased expression for glob 1 (3.8 fold) and Adh (1.5 fold) and decreased expression for pglym78 (0.45 fold), mRpL2 (0.55 fold), obp44a (0.05 fold), retinin (0.031 fold) found as compared to *OregonR^+^* (Figure 5B and 5C).

### Over-Expression of Tau Alters the Mitochondrial Morphology and Membrane Potential

Healthy mitochondrial morphology along with excellent membrane potential was exhibited by the wild type eye imaginal discs as revealed by Mito-GFP and Mitotracker, respectively (Figure 6A-D). The AD eye imaginal discs showed evidence of mitochondrial fragmentation as divulged by damaged mitochondria and leakage of Mitotracker in Cytosol region as a clear evidence for loss of mitochondrial membrane potential (Figure 6E-H). The diseased mitochondria show approximately four fold mitochondrial fragmentation in contrast to control (Figure 6I). The control tissues show two fold mitochondrial elongation in comparison to AD tissues (Figure 6J).

**Figure 6:**
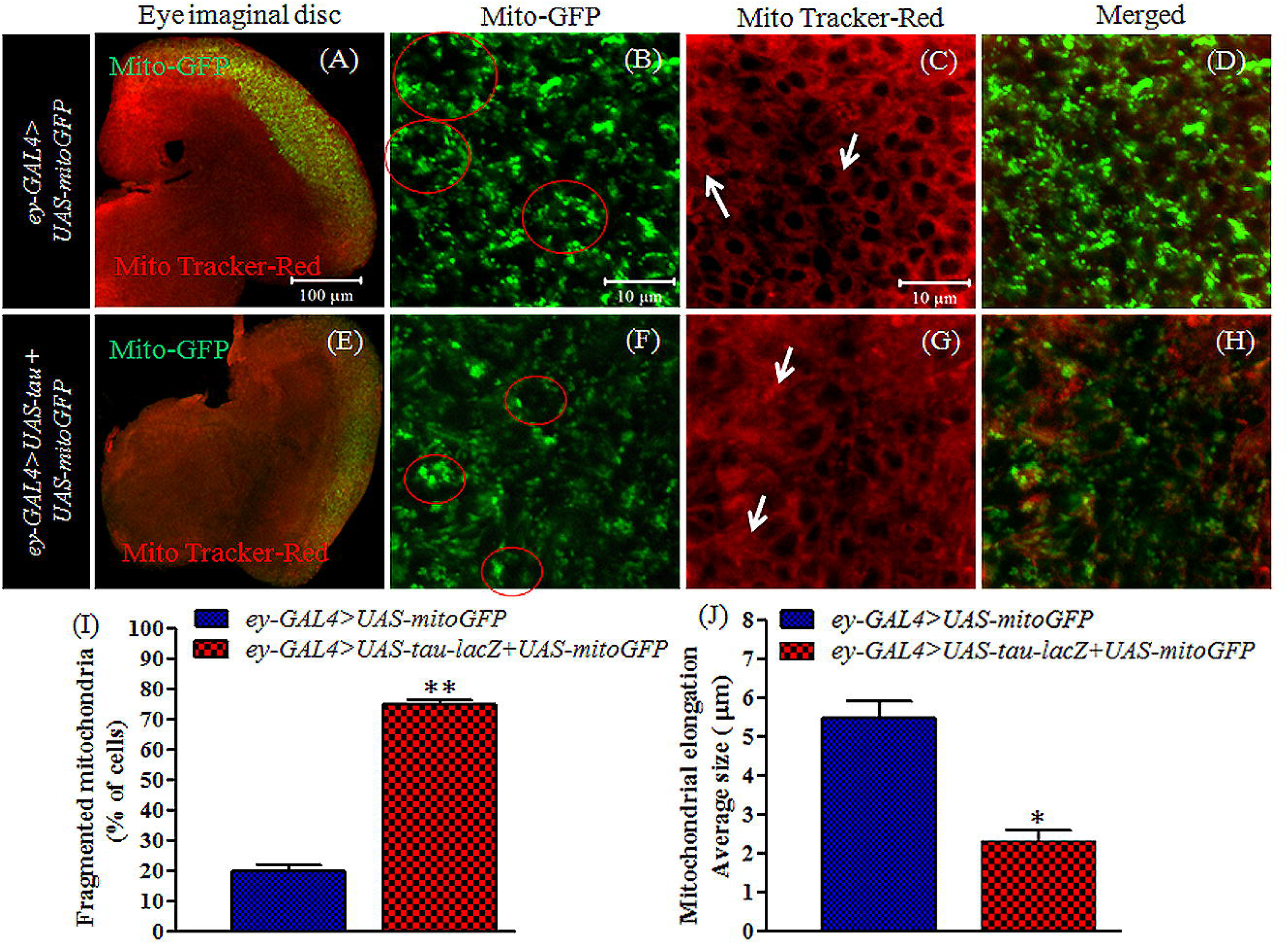
Alteration in mitochondrial morphology in tau over-expressed eye imaginal discs of *Drosophila* 3^rd^ instar larvae as compared to control. The image (A) and (E) represents morphology of eye imaginal disc stained with Mito Tracker-Red as well as express Mito-GFP in control and diseased 3^rd^ instar larvae. Magnified image (B) shows healthy mitochondria represented with Mito-GFP reporter (See red circle). Image (C) also shows healthy mitochondria revealed by Mitotracker accumulation between outer and inner membranes and (D) overlaid image of (B) and (C). (F) Shows fragmented mitochondria (See red circle) and (G) displays damaged mitochondria as revealed by dispersion of Mitotracker (See white arrows). (H) Merged image of (F) and (G). (I) Represent percent cells of fragmented mitochondria in eye imaginal disc of control and AD. (J) Shows measurement of elongated mitochondria in (μm) with the aid of offline software ImageJ 1.38x. The data were analyzed using student’s t test *\*\*p<0.0021*, *\*p<0.0112*.

### Over-Expression of Tau Alters the Mitochondrial Dynamics and Energy Currency

The relative mRNA expression levels of mitochondrial dynamics regulators *Drp-1, Marf-1* and *Parkin* were examined using the head tissues of tau induced AD flies as compared to *OregonR^+^*. The mitochondrial dynamics regulator-1 (*Drp-1*) required for mitochondrial fission exhibits 4.75 fold overexpression (Figure 7A), mitochondrial assembly regulatory factor-1 (*Marf-1*) and *parkin* (ubiquitin ligase) required for mitochondrial fusion displayed more than 2 fold up-expression in the head tissues of AD flies as compared to *OregonR^+^* (Figure 7A). Arguably, the higher degree of *Drp-1* expression as compared to *Marf-1 and Parkin* could cause the elevation of mitochondrial fission than fusion in tau induced AD tissues. In addition, relative expression of HtrA1/2 (mitochondrial serine protease) involved in caspase-dependent apoptosis was also examined and found to exhibit more than 1.5 fold over expression in AD tissues as compared to *OregonR^+^* (Figure 7A). Thus, the mitochondrial dynamic marker studies revealed the basis for the mitochondrial dysfunction in the tau induced AD tissues.

**Figure 7:**
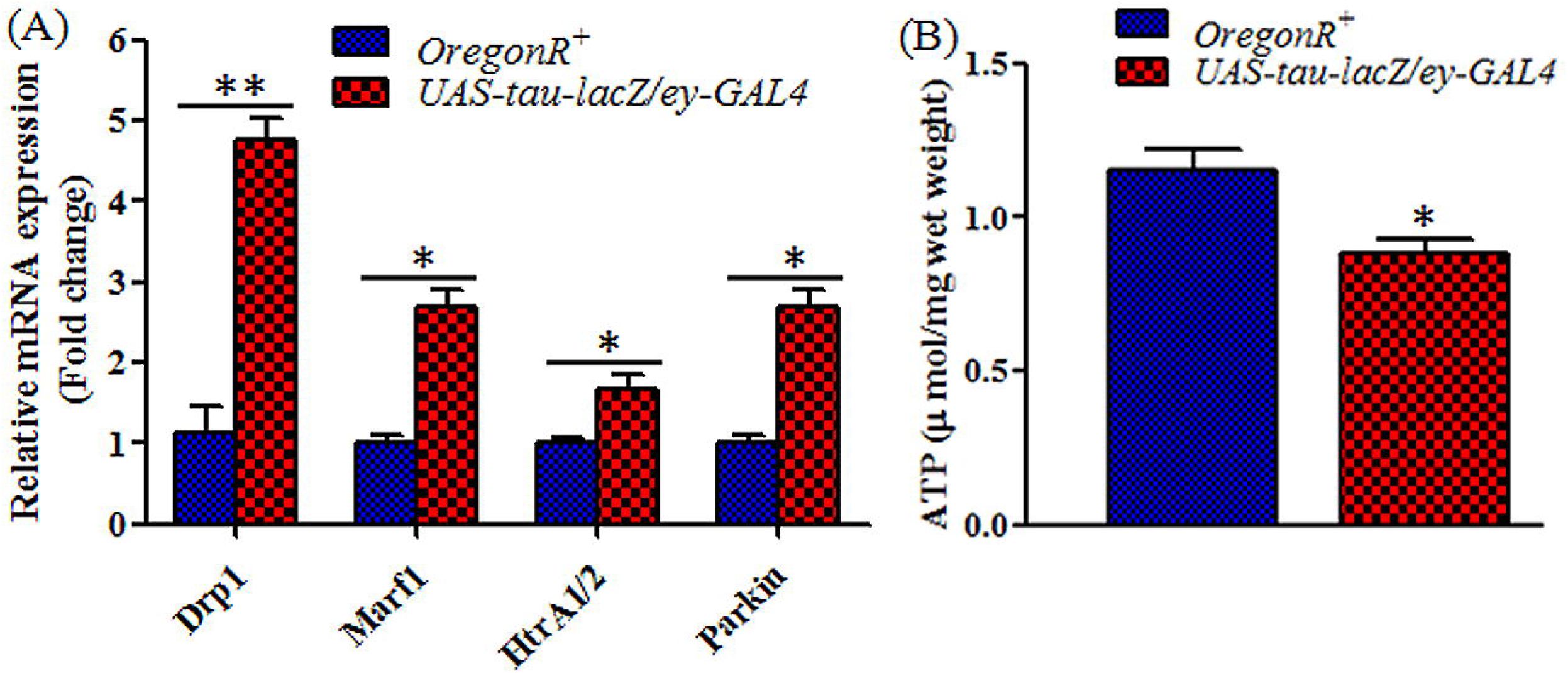
(A) Relative mRNA expression level of mitochondrial dynamics regulators in the head tissues of tau induced AD flies compared to control flies. The data shows significance difference between the two groups using student’s t test *\*p<0.0270*, *\*\*p<0.0015*. (B) ATP quantitation in the head tissues of AD and control flies. The data shows significance difference *\*p<0.0113* using student’s t test in AD group as compared to Control.

Adenosine triphosphate (ATP) is primary energy currency for all living system. ATP quantitation analysis revealed 0.7 fold down regulation of energy currency in the head tissues of AD flies as compared to control (Figure 7B). The mitochondrial impairment could lead to a decrease in ATP level in AD. The newly identified protein, mitochondrial ribosomal protein L2 (mRpL2) that show dysregulation in AD tissues might interact with tumor suppressor protein p53 through novel nucleolar protein 1(Non1) as revealed by flybase interaction tool (Figure 11D). It may be noted that more than 10 fold upregulation of p53 occurs in the head tissue of AD flies as observed by qRT-PCR. The alteration in mitochondrial protein could lead to enhanced cell death in neuronal tissues due to the elevated levels of p53.

### The AD Model of *Drosophila* Exhibits Weight Loss and Less Food Intake

AD flies exhibit loss in body weight compared to controls. This could be due to reduced expression of Odorant binding protein (Obp44a) in olfactory sensilla of tau induced AD flies which could affect the food consumption leading to reduction in body size and weight which is clearly visible in AD flies (Figure 8A, B).

**Figure 8:**
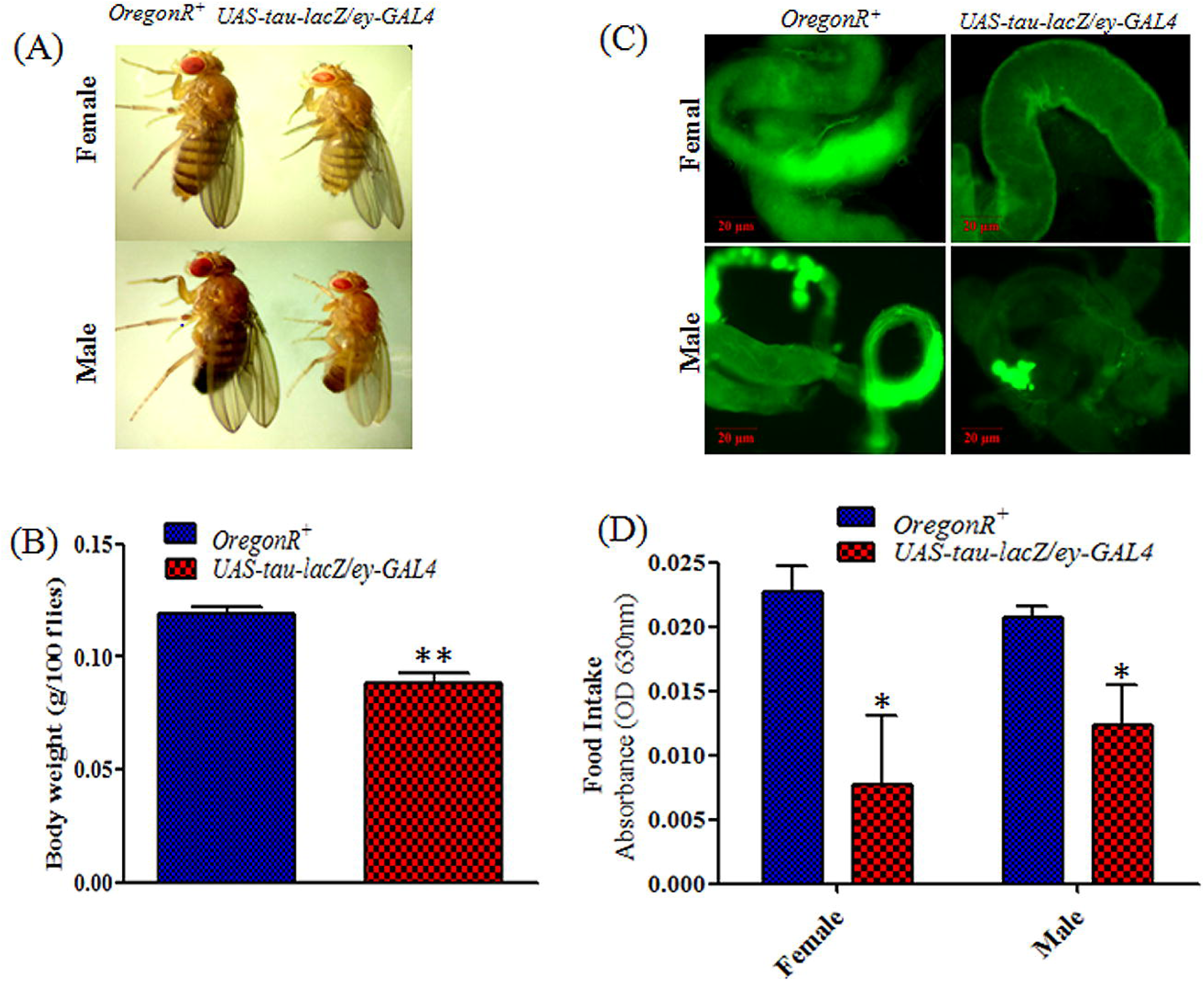
Food intake and body weight measurement of *OregonR^+^* vs. *UAS-tau-lacZ/ey-GAL4.* (A) Displays difference in body size and structural changes of age matched AD flies in contrast to control, (B) Illustrates average body weight of age matched AD flies as compared to control using student’s t test *\*\*p<0.0040*, (C) food tracing dye in gut tissues of respective flies, (D) Illustrates absorbance of supplemented food containing tracing dye in gut tissues. The data shows significant difference *\*p<0.0221* in AD group as compare to control using student’s t test.

**Figure 9:**
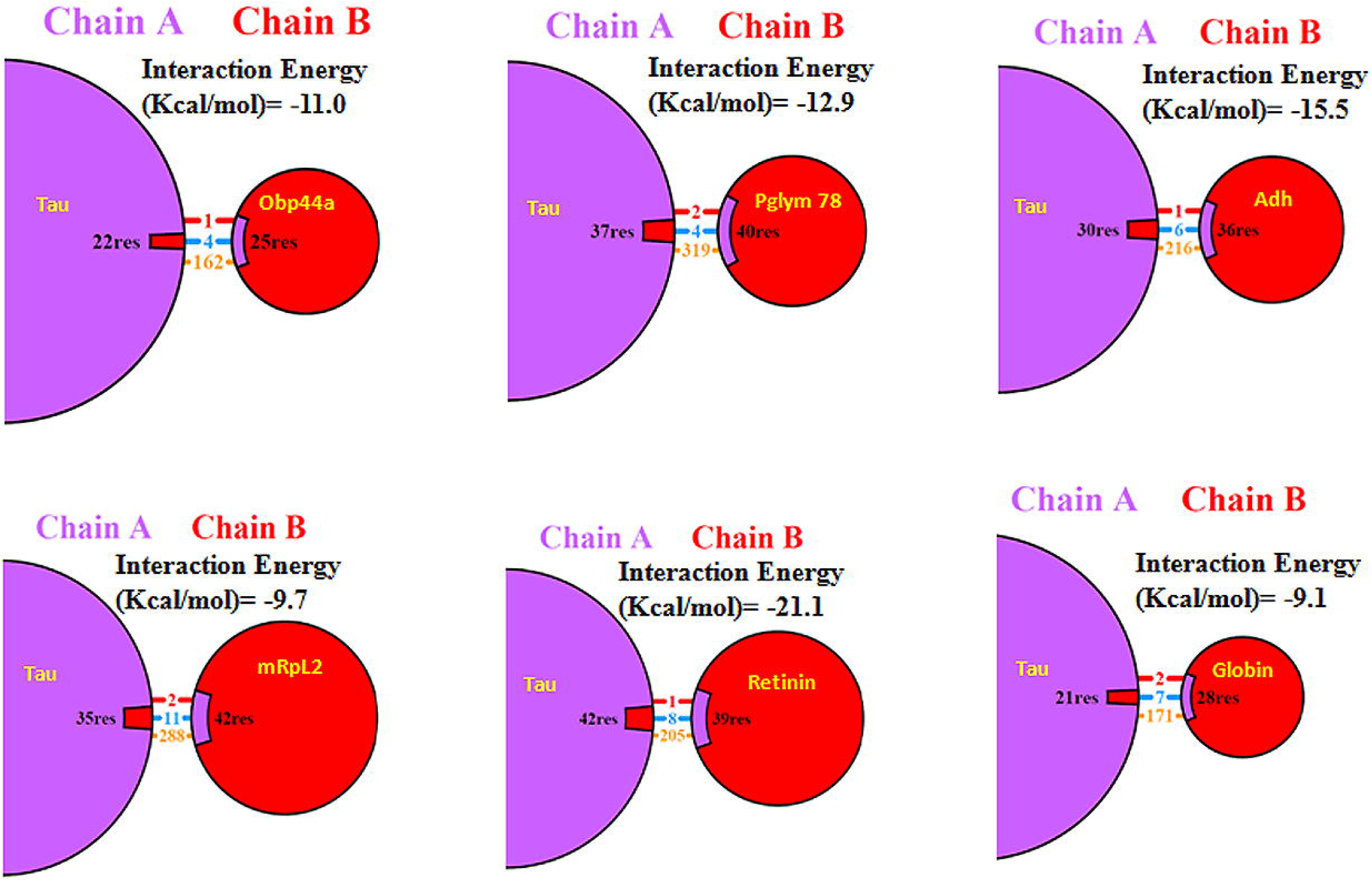
Summary of the interactions of differential expressed identified Proteins (Chain B) with Tau Protein (Chain A) of *Drosophila melanogaster.* Interaction energy (Kcal/mol) and Interaction Colour Code-Red: Salt Bridge; Blue: Hydrogen Bonding; Orange: Non-bonded contact.

**Figure 10:**
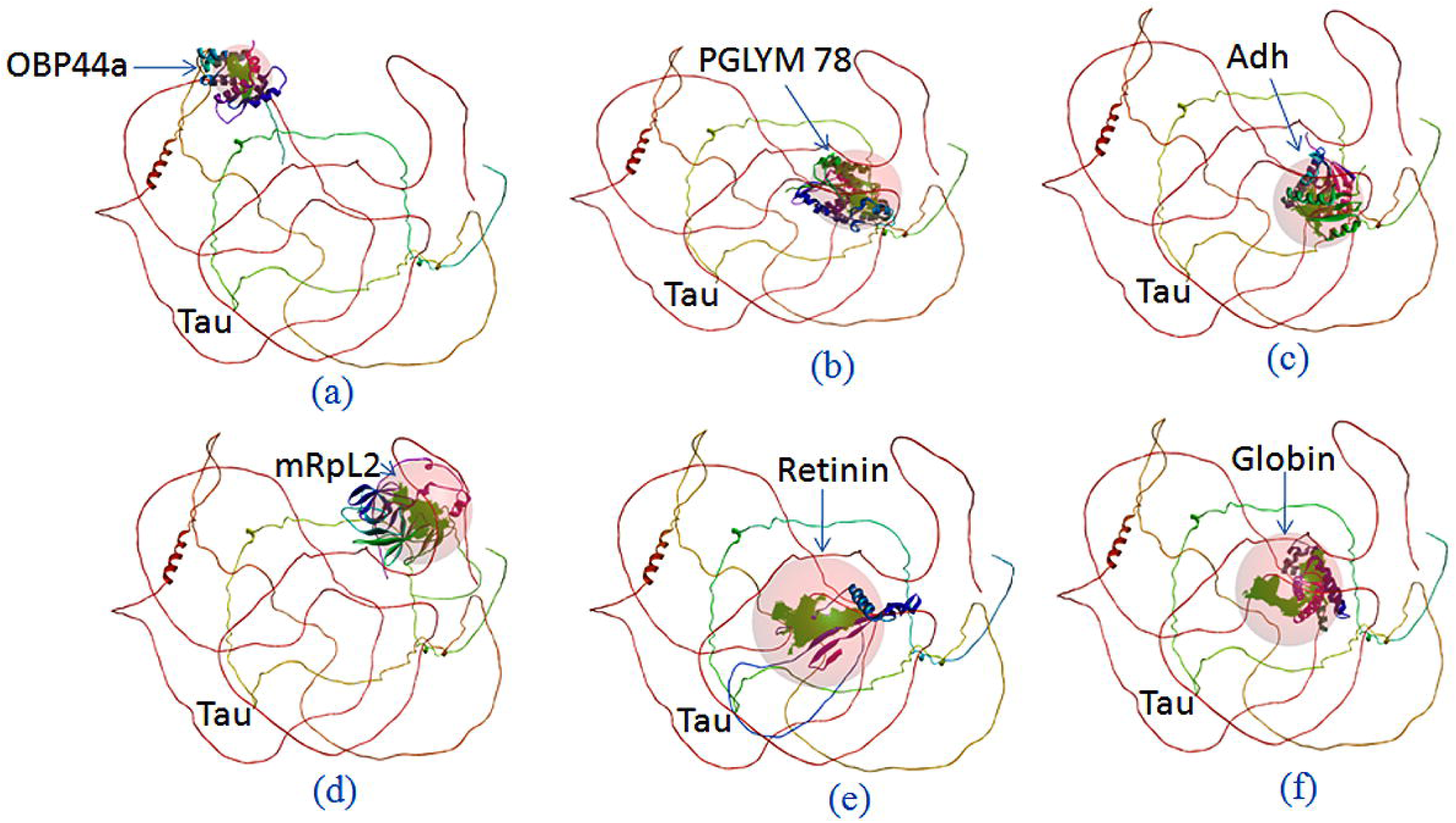
Representation of the identification of Tau pocket to host identified proteins of *Drosophila melanogaster*.

Hence, to examine the possible cause of body weight reduction in AD model of *Drosophila*, the food intake measurement analysis was also carried out using liquid synthetic food tracer green dye (Manufactured by TRISHUL group). This assay revealed that tau induced AD flies exhibited less food intake as compared to control as shown by lesser accumulation of colored food in gut tissues of the flies (Figure 8C). The absorbance of fed food in gut tissues of diseased flies was monitored and found to display reduced absorbance as compared to *OregonR^+^*(Figure 8D).

### Cell Viability and Mortality Assay

The metabolic activity was examined using 3^rd^ instar larvae of *OregonR^+^ and UAS-tau-lacZ/ey- GAL4-UAS-*GFP strain (Figure S4A). The eye imaginal discs were dissected out for measuring the cell viability. In cell viability experiment, yellow tetrazolium salt reduces to form insoluble formazan crystal through the action of cytosolic enzyme, NAD(P)H-dependent oxido-reductase, present in active cells (Berridge et al., 2005). The *OregonR^+^* eye imaginal discs exhibited greater metabolic activity, showing higher enzymatic (Oxidoreductase) activity, leading to increased MTT reduction. These findings indicated that AD eye tissues exhibited reduced metabolic activity in contrast to control (Figure S4B). Moreover, the survival curve showed that control flies (*OregonR*^+^ and *elavc155-GAL4*) show extended life span as compared to tau induced AD model of *Drosophila.* The control flies exhibited about 1.3 fold higher life spans in comparison to diseased flies (Figure S4C). Maximum life span was observed in *OregonR^+^* flies. The mortality assay was repeated thrice in each category of flies. The median survival for *OregonR^+^, elavc155-GAL4* and *elavc155-GAL4>UAS-tau-lacZ* were 70, 67 and 44 days respectively (Figure S4D). The Log-rank (Mantel-Cox) test was applied to perform the significance analysis of survival data. The results showed a significance difference between controls and AD flies reared in normal food \*\*\**p<0.0001*.

### Olfaction Analysis in Tau flies

The Y maze assay that represents a well-organized test to assess the chemosensory response was performed to examine sensory perception of olfactory organ in *OregonR^+^* and AD flies. The study revealed that *OregonR^+^*flies exhibited higher attraction towards Acetic acid (AA) than the AD flies. AD flies were visible in very small numbers than the control flies towards the odorant molecule (Figure S3).

### Protein-Protein Docking Analysis

The protein-protein docking study of Tau protein with newly identified proteins from MALDI-TOF/MS were analyzed in order to obtain the interactions energy and bond interactions between proteins by accounting for steric fit and chemical complementation. Herein, the interaction energy of Tau protein with OBP44a, PGLYM78, Adh, mRpL2, retinin and Globin were scored to be -11.0, -12.9, -15.5, -9.7, -21.1 and -9.1 kcal/mol, respectively. The protein-protein docking study predicted that tau protein has strong affinity with Retinin followed by Adh, Pglym78, Obp44a, mRpL2, and Globin. Tau-Retinin established better interactions energy (kcal/mol) because of polar-polar interactions unlike the other identified proteins. However, different bond type interactions were observed between the Tau protein and identified proteins such as hydrogen bondings, salt bridges and non-bonded interactions. The numbers of interchain hydrogen and salt bridge bonds flanked by Tau protein with identified proteins such as OBP44a, Pglym78, Adh, mRpL2, retinin and Globin were (4,1), (4, 2), (6, 1), (11, 2), (8,1) and (7, 2), respectively was observed while rest of the interactions were non-bonded contacts. In such condition, mRpL2 exhibited the maximum numbers of nonpolar interactions with polar amino acids of tau protein. In case of Tau-Globin, very poor interaction has been seen which could be due to rich nonpolar-nonpolar interactions between amino acid residues (Figure 9). Further, the details of hydrogen and salt bridge interactions at amino acid residues level with distances between Tau and associated identified proteins have been shown in Table S1. In addition, pictorial representation of Docked protein’s surface structures and hydrogen bond interactions of these novel proteins with tau were analyzed and identified a unique Tau pocket to the host identified proteins have been depicted in Table S2, Figure 10. Moreover, different bond type interactions such as hydrogen bonds, salt bridges, disulphide bonds and non-bonded contacts of tau protein with identified proteins and the corresponding Ramachandran Plots for the interactions of the same have been shown in Table S3.The construction of Ramachandran plot permitted the complex structures of docked proteins used to determine main secondary structure elements (α-helix and β-sheet) clustered within the allowed and disallowed regions (Table S3). Ramachandran plot aid in predicting the descriptive conformation of amino acid residues of docked proteins of Tau with OBP44a, Adh, Pglym78, mRpL2, Retinin and Globin proteins were found be 48.1, 54.3, 54.2, 53.0, 47.6, 50.9% in favoured region and (26.1, 9.6%), (23.1, 8.2%), (23.3, 8.1%), (23.7, 8.7%), (26.2, 10.3%), (23.9, 9.1%) in additional and generously allowed regions and rest amino acid residues were found to be 16.2, 14.4, 14.4, 14.6, 15.8, and 16.1% in disallowed regions (Table S3).

### In Silico Interaction Analysis of Identified Proteins

The network interaction depicts the association of differentially identified proteins with other proteins. The network interactions of the identified proteins were explored via online tools “fly base interaction browser” and “esyN network builder”. The Odorant-binding protein (Obp44a) shows common association with Apolipophorin (Apo1pp) and Odorant binding protein 56d (Obp56d). The Obp44a also indirectly linked with Phosphoglyceromutase (Pglym78), Alcohol dehydrogenase (Adh), and Wingless (wg) through Apo1pp. Moreover, Obp44a exhibit interaction with Actin (Act79B) cytoskeleton protein and adult cuticle protein 1 (Acp1) via Obp56d. This suggests that Apo1pp and Obp56d are crucial proteins exhibits interaction with other proteins to exert the chemosensory function (Figure 11A). Similarly, Pglym78 shows direct relationship with Adh, Apo1pp, Enolase (Eno), Rab11 and CG2493 (Figure 11B). The Adh also directly linked with Pglym78 and Eno (Figure 11C).

**Figure 11:**
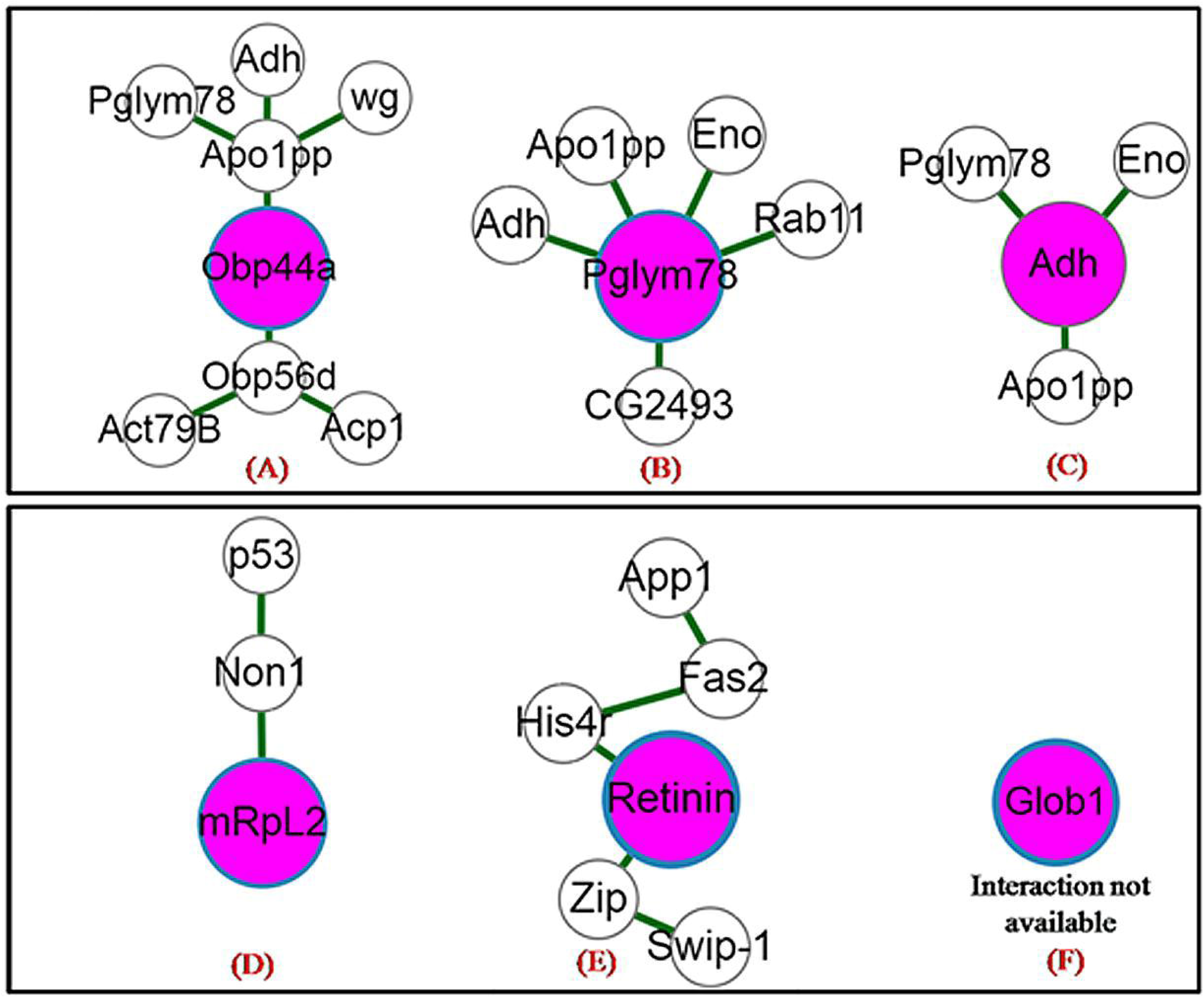
Flybase interaction browser depicts the regulation of other proteins through differentially identified proteins. The network interaction of individual identified protein was build via online esyN network builder. The purple colour balls representing the individual protein identified from MALDI-TOF/MS analysis. The identified protein in image (A) Obp44a, (B) Pglym78, (C) Adh, (D) mRpL2, (E) Retinin, and (F) Globin1 exhibited directly or indirectly regulation of other proteins.

On the other hand, mitochondrial ribosomal protein L2 (mRpL2) linked with tumor suppressor protein p53 through Novel nucleolar protein 1(Non1) (Figure 11D). Further, the retinin protein interconnected with Fasciclin 2 (Fas2) through Histone H4 replacement protein (Figure 11E). The other identified protein Glob1 do not showing any interaction with proteins present in fly base data (Figure 11F).

### Biocomputational Classification and Pathway Analysis of Identified Proteins

The distribution of six identified proteins was classified according to molecular function, biological process, cellular components, and protein classes in percentage using online panther database. In molecular function, Adh protein exhibited 50.0% catalytic activity and mRpL2 displayed 50.0% structural molecule activity (Figure S5A).This outcome predicted that Adh may involve in Alcohol metabolism and produces more toxic acetaldehyde during AD progression. In biological processes, mRpL2 display only 33.3% role in metabolic process means body gets very less energy through oxidative process from food in disease condition and 33.3% in cellular process, while Obp44a involved in 33.3% multicellular organismal process (Figure S5B). In cellular components majority of proteins like mRpL2 and Adh together shares 40.0% cell part. The signaling molecule Obp44a represent 33.3% part involved in extracellular region and mRpL2 existed as the 33.3% part of macromolecular complex (Figure S5C). In protein classes, mRpL2 shows 33.3% nucleic acid binding protein, Adh represents 33.3% Oxidoreductase activity and Obp44a exhibited as 33.3% signaling molecule shown in following (Figure S5D).

Further, analysis of Panther pathway predicted the involvement of identified protein Phosphoglycerate mutase (Pglym78) in metabolic process like glucose metabolism. The reports suggested that Pglym78 also involved in several pathways like oxidative stress, insulin signaling and neurodegeneration diseases. Thus, study concluded that differentially identified proteins through proteomic approach play crucial role and exhibited it’s up and down regulation in AD.

### The Identified Proteins from MALDI-TOF Reflects Evolutionary Relationship

The phylogenetic tree has been drawn in blocky format. The optimal tree with the sum of branch length = 7.11589613 has been shown. The phylogenetic tree revealed a better comparison of protein sequences and evolution at the molecular level. Similar protein sequences of proteins exhibited very close evolutionary relationship. For instance, the protein retinin with tau, Obp44a with globin1 isoform B, and IP15846p (Adh) with RE45450p (mRpL2) infer the close evolutionary relationships and also represented equal evolutionary distance of branches to reconstruct ancestral states of proteins (Figure S6). Hence, using this information, we can trace the paths followed by evolution. Those proteins which show large compositional variation represented less relationship. Thus, the branching pattern in evolutionary tree clearly indicates the close relationship of proteins with each other.

## DISCUSSION

Currently not many reports are available on Tau specific proteome analysis in relation to the AD. In order to understand Tau specific proteome and its role in the disease progression, we have for the first time identified a tau induced proteome through AD model of *Drosophila.* Further, the newly identified proteome candidates were analyzed for Tau specific interaction by *in silico* methods. A previous study had revealed that upregulation of p53 gene in AD induces the tau phosphorylation in cells (Hooper et al., 2007; Chang et al., 2012). In present study also we observed that the p53 expression was elevated to over 10 fold in AD flies, in contrast to control, in agreement with the previous observations. Elevated levels of tau and p53 could be responsible for the death of ommatidia and reduction in eye size (Figure 1E, F). The 2D proteomic study was done to identify the novel tau associated proteins. The MALDI-TOF/MS analysis revealed six proteins Obp44a Isoform A, Pglym Isoform A, Adh variant, mRpL2, Retinin, and Glob1 Isoform B as differentially expressed under conditions of Tau induced AD. There are few reports available for role of Alcohol dehydrogenase and the brain Globin in AD pathogenesis (Balcz et al., 2001; Xie and Yang, 2016). It has also been documented that upregulation of p53 suppress the Phosphoglycerate mutase (Pglym) activity through which it induces AD progression (Liu and Xu, 2011). According to recent reports, upregulation of p53 enhances the tau mediated AD pathogenesis (Jembrek et al., 2019). The present study also observed similar activity in tau induced AD model of *Drosophila* (Figure1S). Till date there are no studies implicated on role of Retinin protein in tauopathies. However, several reports suggest that tauopathies could result due to mitochondrial dysfunction, oxidative stress, weight loss and dietary dysfunction, etc (Sultana et al., 2007; Smith and Greenwood, 2008; Moreira et al., 2010; Kai et al., 2015). In light of this, the mRpL2 and Obp44a proteins emerged as potential candidates to work on mechanistic pathways connecting the mitochondrial biogenesis activities. The possible relationships of differentially expressed proteins (mRpL2 and Obp44a) with hyperphosphorylated tau protein have been shown in following proposed AD pathway (Figure S7).

Herein, we showed for the first time that over expression of tau altered the expression of mRpL2 and Obp44a during AD progression in *Drosophila*, although exact mechanism is still elusive. Recent studies reported that hyper phosphorylated tau protein perturbed the mitochondrial dynamics via failure of the axonal transport and loss of synaptic function (Cai, 2017; Cheng and Bai, 2018). Loss in axonal transport of mitochondria elevates the dissipation of energy currency in the brain tissues through which neuroinflammation and oxidative stress are generated (Errea et al., 2015). Another study revealed that altered expression in mitochondrial ribosome protein (MRP) enhances the neurodegeneration (Lunnon et al., 2017). In the present study the reduced ATP levels in AD head tissues could be due to reduced expression of mRpL2 (Figure 5A, 7B). In turn, the down-regulation of mRpL2 in the head tissue of AD flies could be responsible for generation of reactive oxygen species (ROS) and enhance dissipation of energy currency. Thus, these findings suggested that over-expression of Tau protein in eye imaginal disc of *Drosophila* 3^rd^ instar larvae can cause reduced expression of mRpL2, which would be responsible for alteration in mitochondrial structure as well as mitochondrial membrane potential. The impairment in mitochondrial morphology and membrane potential could be consequence of tau hyperphosphorylation. The novel mitochondrial ribosomal protein L2 (mRpL2) normally assist in protein synthesis within mitochondria; while it exhibited reduced expression in AD in contrast to control (Figure 5A). The tau induced AD tissues illustrate discrepancy in mitochondria morphology and loss in membrane potential in contrast to control using *UAS-Mito-GFP* and Mito-Tracker Red also in agreement with these observations (Figure 6A-H).

Further, the enhancement of mitochondrial fragmentation in the eye tissues of AD flies also could be due to alteration in the expression of mitochondrial dynamics regulators such as *Drp1, Marf1, HtrA1/2, and parkin* (Figure 6F, 7A). The key function of mitochondrial dynamic regulators like *Drp1* in hyperphosphorylated tau induced AD has been widely explored (Manczak and Reddy, 2012). A previous study reported that imbalance of mitochondrial distribution impaired their fission and fusion process in AD (Wang et al., 2009; Zhu et al., 2013). This process could be due to reduced expression of mRpL2 in the head tissues of AD flies. Earlier studies also revealed that metabolic dysfunction is a consequence of AD progression (Cai et al., 2012), and phosphorylated tau induces anorexia (loss of appetite) due to alteration in olfactory proteins making the food less appealing resulting in loss in body weight (Gillette-Guyonnet et al., 2007; Tamura et al., 2008; Ansoleaga et al., 2013; Stamps et al., 2013; Yoo et al., 2017; Martinez et al., 2018). Further, chemosensory proteins are required for sensory perception of chemicals and food detection (Shanbhag et al., 2001; Pinto, 2011; Martinez et al., 2018). In present study, the reduced expression of chemosensory protein, Obp44a, enhances mitochondrial stress, causing the nutritional anxiety reflected as symptom of AD (Figure 8A-D). Hence, alteration in olfactory sensillary proteins such as Odorant binding protein 44 a (Obp44a) could be responsible for development of anorexia symptoms which ultimately cause loss of appetite manifested in AD flies. These consequences might be due to very complex network of optic neurons to olfactory neurons in AD fly heads. The Tau induced deterioration in compound eyes of AD flies enhance the degeneration of optic neurons through which it impedes the consistency of associated network of olfactory neurons (Figure 2I, II) leading to the alteration in expression of Obp mRNA in olfactory sensilla.

Further, cell viability assay was performed to perceive the viable cells in eye imaginal tissues. The diseased imaginal discs exhibited reduced cell viability in contrast to control (Figure S4B). The loss in cell viability could be reason of nutritional stress which ultimately affects the various cellular functions.

The earlier reports revealed that mutation in mitochondrial ribosomal protein (MRP) in C. elegans reduces their life span (Mozhui et al., 2017). The tau induced AD model of *Drosophila* also exhibited reduced life span, which could be due to altered expression of mRpL2.The *in-silico* studies revealed that the newly identified proteins exhibit direct or indirect interactions with other proteins implicated in AD, which are governed through different regulatory pathways (Figure 11A-F). One of the identified proteins, Pglym78 directly associated with Adh, Eno, Rab11 (Figure 11B). The normal function of Eno involved in glucose metabolism but alteration in its expression contributed dementing disorder progression (Butlerfield and Lange, 2009). Alternatively, Rab11 protein has been implicated in recycling of β-secretase to the plasma membrane. The presence of β-secretase enhances proteolytic activity for the Aβ production (Bhuin and Roy, 2015). Moreover, mRpL2 directly linked with p53 protein and involved in mitochondrial translation in neuronal cells, while p53 protein induces neuronal cell death phenomenon. Thus, the elevated expression of p53 protein and loss of mRpL2 in present study is in agreement with earlier reports showing involvement of p53 protein in dead and damaged neurons (Morrison and Kinoshita, 2000). Similarly the involvement of cornea-specific protein Retinin linked with Fas2 via core Histone protein H4. The altered expression of Retinin in the degenerated ommatidia of reduced eyes of AD flies is an indicative of Tau mediated phenomenon (Figure 5). However, the exact role of Retinin protein is not known well and it may be participated in optical property or refractive power like to drosocrystallin, a glycoprotein of the *Drosophila* corneal lens (Kim et al., 2008). Fas2 facilitates neuronal cell adhesion and regulation of synaptic growth at neuromuscular junction. Mis-expression of Fas2 protein in photoreceptors causes neuronal targeting defects.

The biocomputational classification of novel proteins shows a number of molecular, biological, cellular and protein class activity as revealed by online panther database (Figure S5A-D). Interestingly, the mRpL2 protein participates in good proportion of metabolic activities as revealed by biological processes (Figure S5B). The protein-protein docking studies of Tau protein with mRpL2 exhibited weak interaction energy which could be due to nonpolar-polar interactions of amino acids, while Globin exhibited very poor interaction energy with Tau protein as revealed by richness of nonpolar-nonpolar amino acid interactions (Figure 9). Interestingly, Retinin depicting strong interactions energy with Tau protein because of sharing more polar-polar amino acids between both species, however other identified proteins Adh, Pglym78 and Obp44a exhibited the descending order of interaction energy (Figure 9). Interestingly, the interaction of Retinin, Adh, Pglym78 with Tau proteins exhibited similar binding cleft unlike others proteins. This Tau pockets cleft rich in acidic and basic amino acids sharing with the host identified proteins could be key target for future studies (Figure 10, Table S2). The altered expression of mRpL2 protein enhances the oxidative stress production in mitochondria. The oxidative stress production contributes significant role in p53 over expression, ATP dissipation etc. Thus, altered levels of p53 and newly identified proteins probably participate in aggregation of phosphorylated tau protein towards NFT formation, which ultimately enhances AD pathogenesis. Thus, further studies are needed for finding the functional relationship of newly identified proteins with tau protein to investigate the exact route of tau mediated AD pathogenesis.

## CONCLUSION

Neuroscientists are investigating the tau associated route of AD progression for the last several decades. Till date no clear evidence of molecular basis of Tau mediated AD pathogenesis has been explored. Herein, for the first time, the eye tissue specific over expression of tau induced proteome analysis was done in *Drosophila* to unravel novel protein candidates through proteomic and insilico approaches in tauopathy flies. The over expression of tau revealed altered expression of several proteins, which can be potential candidates to understand mechanism of AD progression. Of these, six proteins viz., Obp44a, Pglym78, Adh, mRpL2, and Retinin proteins were characterized by proteome and *in-silico* analyses. The altered mitochondrial morphology, membrane potential and loss in body weight of AD flies could be due to down regulation of mRpL2 and chemosensory Obp44a protein while eye degeneration phenotypes could be linked to altered expression of retinin protein. The impairment in mitochondrial dynamic regulators, mitochondrial membrane potential, energy currency, food intake, and body weight potentially triggers the AD pathogenesis. In end, *insilico* interaction study of tau protein with newly identified altered proteins from the host strongly implicates their role in pathophysiology of AD and could be explored for possible therapeutic targets against Tauopathies.

## AUTHOR CONTRIBUTIONS

BSC designed and performed most of the experiment, analyses the data and wrote the main manuscript. SS involved in all aspects of studies and supervised the whole experiments. JS assisted in reagents preparation for 2D Gel electrophoresis. SG involved in protein-protein docking, refinement and data analysis in this manuscript.

## FUNDING

This work was supported by the IoE, BHU and a research grant (P07/598) awarded to S. Srikrishna by the SERB, New Delhi, India and authors are highly grateful to SERB for Financial support. Authors are thankful to IoE, Banaras Hindu University for intramural grant to SS. Central facility, ISLS, SATHI, Banaras Hindu University are greatly acknowledged. One of the Authors (B.S. Chauhan) is grateful to ICMR (Letter no. 45/50/2018-PHA/BMS) for providing financial support in the form of Senior Research fellowship. The financial support by Institutions of Eminence (IoE), BHU is highly appreciated.

## ACKNOWLEDGEMENT

The central facility support extended by the DST-FIST Level-II, Biochemistry department (SR/FST/LS-II/2021 - TPN - 70458) is acknowledged. We would like to acknowledge ISLS, DBT BHU for Proteomics service, Prof. N.V. Chalapathi Rao for providing SEM facility at Department of Geology, BHU, Varanasi and Prof. S. C. Lakhotia for providing DST-National Confocal Facility at Department of Zoology, BHU, Varanasi, India. We are also highly thankful to Dr. Andrea Brand, University of Cambridge for providing us UAS-tau-lacZ fly stock. We are also highly thankful to the DST-SATHI and ISLS for central facility.

## CONFLICT OF INTEREST

The authors declare that there is no any conflict of interest.

## ABBREVIATIONS

2D-GE: Two Dimensional-Gel electrophoresis
AD: Alzheimer’s disease
Adh: Alcohol dehydrogenase
ATP: adenosine triphosphate;
Drp1: Dynamin-related protein-1
glob1: Globin 1
HtrA1/2: mitochondrial serine protease
IEF: Iso-Electric Focusing
MALDI-TOF/MS: Matrix Assisted Laser Desorption Ionization-Time of Flight /Mass Spectrometry
Marf1: Mitochondrial assembly regulatory factor-1
mRpL2: mitochondrial ribosomal protein L2
MRPs: mitochondrial ribosomal proteins
MTT: 3-(4, 5-dimethylthiazol-2yl)-2, 5-diphenyl tetrazolium bromide
NFTs: neurofibrillary tangles
Obp44a: Odorant binding protein 44a
ORNs: olfactory receptor neurons
Pglym78: Phosphoglycerate mutase 78
qRT-PCR: quantitative real time-Polymerase chain reaction
SEM: Scanning electron microscope
Tau: Tubuline associated unit

## Notes

### Competing Interest Statement

The authors have declared no competing interest.

