## Supplementary material for "Identification of differentially expressed novel tau associated proteins in transgenic AD model of *Drosophila* - a proteomic and *in silico* Approach": Supplementary_Information.docx

***Corresponding Author:***

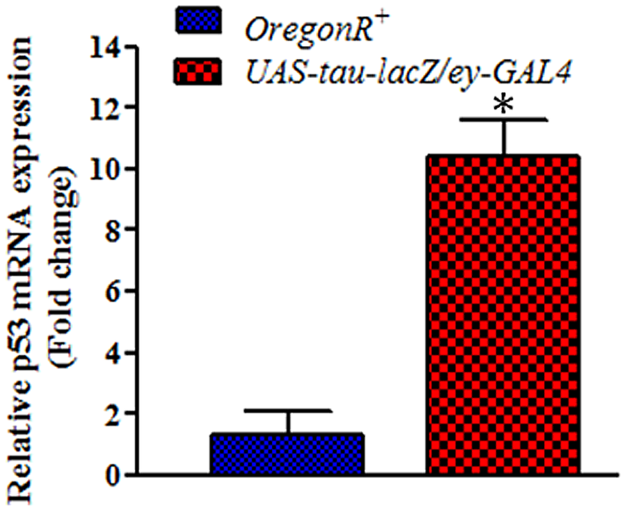

**Figure S1:** Represents the over expression of p53 mRNA in the head tissues of Tau flies in comparison to control with significance difference of **p<0.0236*.

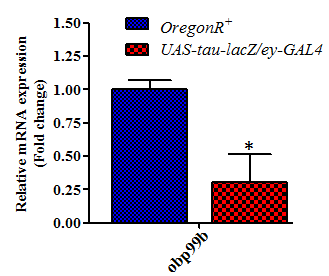

**Figure S2:** Relative mRNA expressions of obp99b gene in head tissues of control (*OregonR^+^*) and experimental sample (*UAS-tau-lacZ/ey-GAL4*). The results showed a significance difference between control and test sample reared in normal food **p<0.0484*.

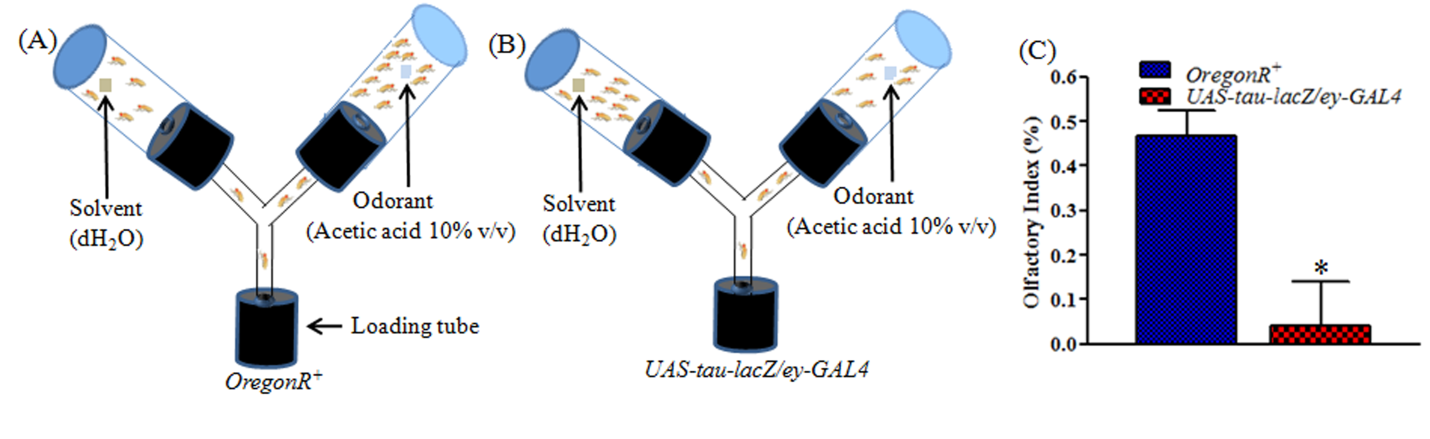

**Figure S3:** Olfactory response of *OregonR^+^* and *UAS-tau-lacZ /ey-GAL4* flies assessed with Y-maze set-up. (A and B) represents the assembled device showed Y-maze assay to evaluate the chemosensory responses in flies.(C) Quantification of Control and AD flies olfactory response towards Acetic acid (AA) diluted in dH_2_O (10% v/v). Total no. of fruit flies for each group (N=150) and t test shows the significance difference **p< 0.0214* in AD group as compared to Control.

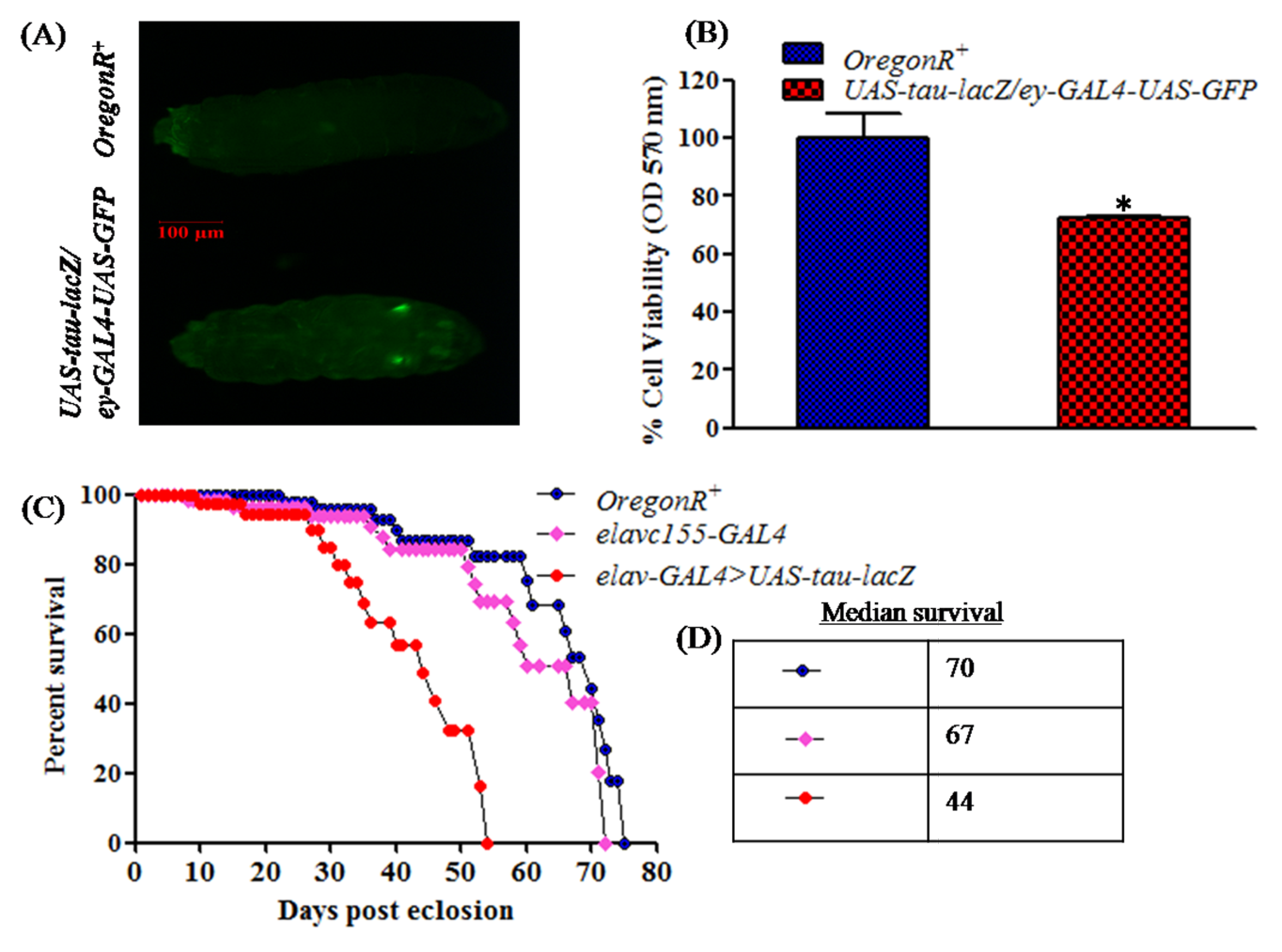

**Figure S4:** The histogram depicted the cell viability and mortality assay of *Drosophila*. The image (A) represents GFP positive screening of transgenic *Drosophila* larvae in contrast to control and histogram (B) exhibits measurement of percent cell viability using student’s t test **p<0.0340*. The curve graph (C) illustrates percent survival of *OregonR^+^*, *elavc155-GAL4*, and *UAS-tau-lacZ* driven with *elavc155-GAL4* driver whereas table (D) displayed their median life span of respective flies stocks.

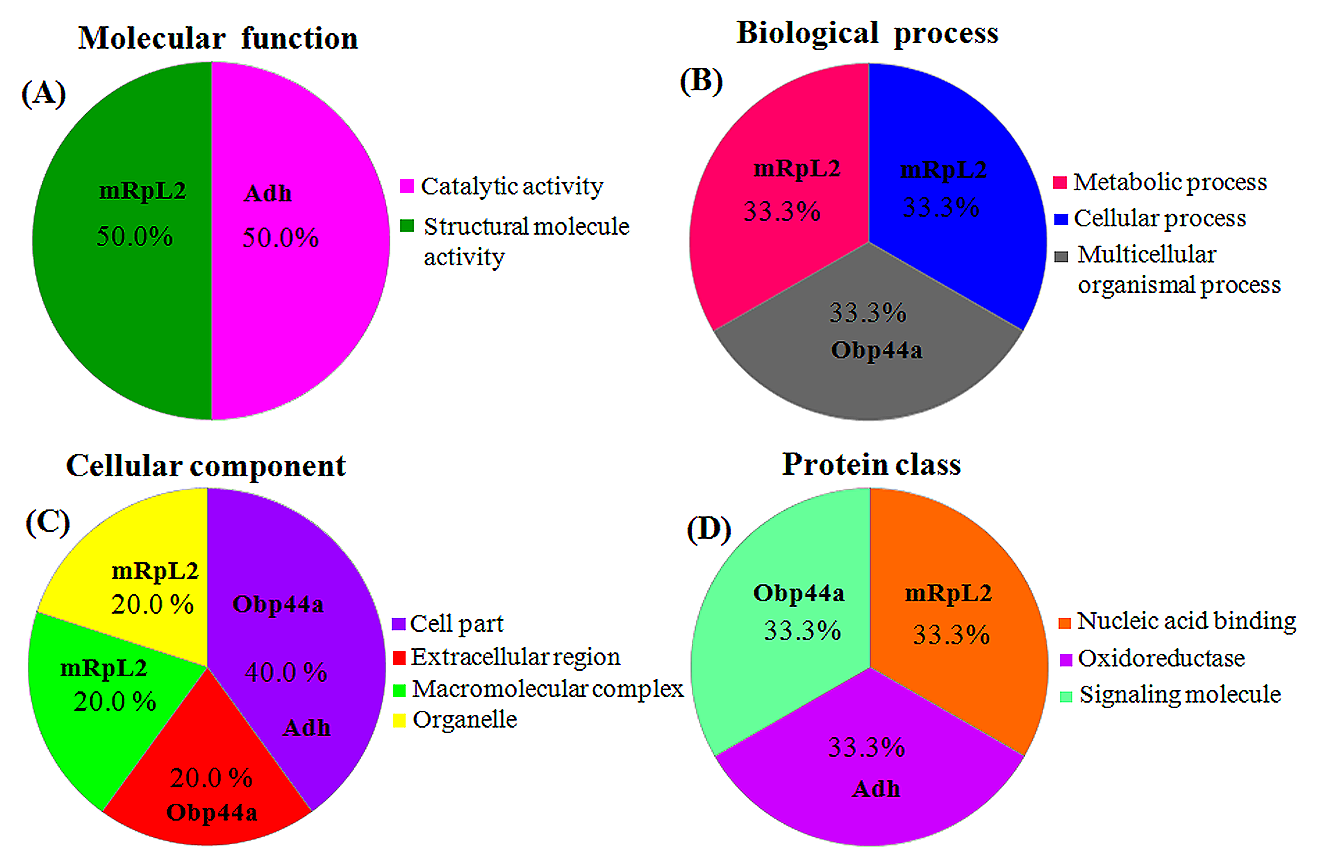

**Figure S5:** Pi-chart illustrated the functional classification of differentially expressed identified proteins in the head tissues of tau induced AD fly. The proteins are classified in following order; (A) molecular function, (B) biological process, (C) cellular component and (D) protein class.

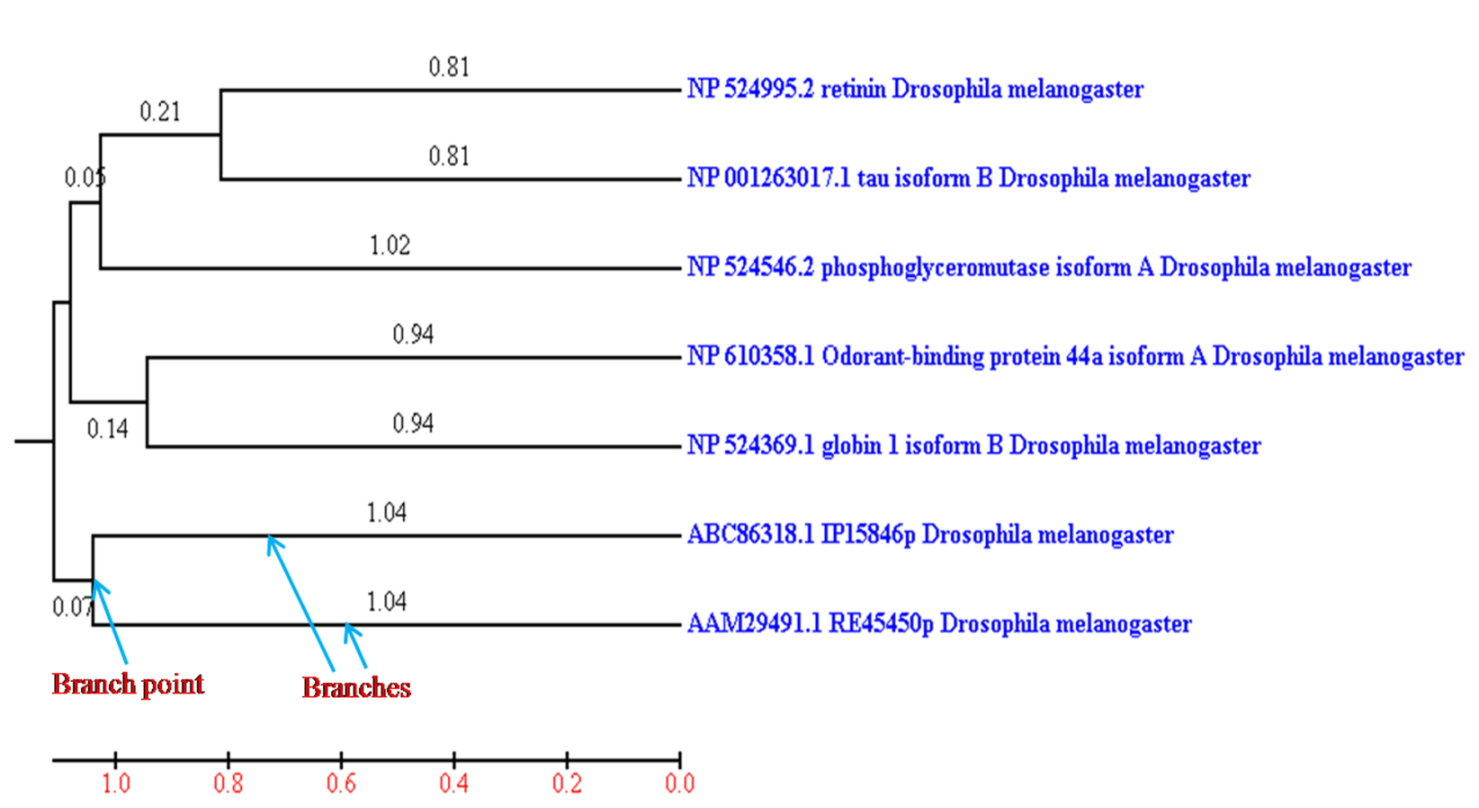

**Figure S6:** Phylogenetic tree reflecting the evolutionary relationship of identified proteins from MALDI-TOF/MS which are positioned at the ends of the branches.

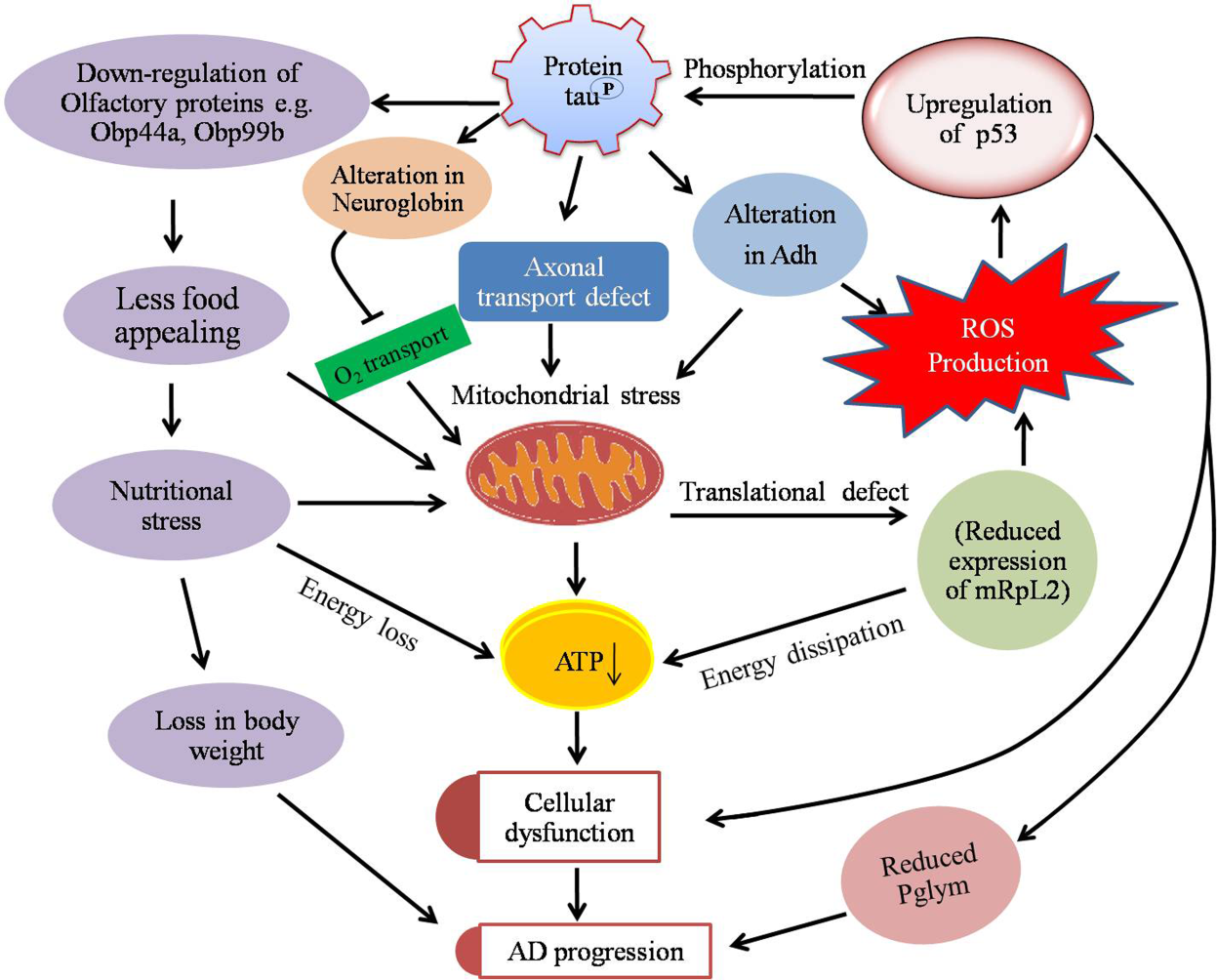

**Figure S7**: The proposed pathway likely represents the initiation of AD pathogenesis. The over expressed tau protein impairs the axonal transport. The alterations in expression of novel proteins lead to elevation of mitochondrial dysfunction as an evidence for AD progression.

**Table S1:** Represents the of the hydrogen and salt bridge interactions at residue levels of identified proteins OBP44a, PGLYM78, ADH, mRpL2, Retinin and Globin (Chain B, Sky) with Tau Protein (Chain A, Green) of *Drosophila melanogaster.*

| **Tau-OBP44a** | | | | | | | | | | | | | | | | | | | | | | | | | | | | | | | | | |
| --- | --- | --- | --- | --- | --- | --- | --- | --- | --- | --- | --- | --- | --- | --- | --- | --- | --- | --- | --- | --- | --- | --- | --- | --- | --- | --- | --- | --- | --- | --- | --- | --- | --- |
| 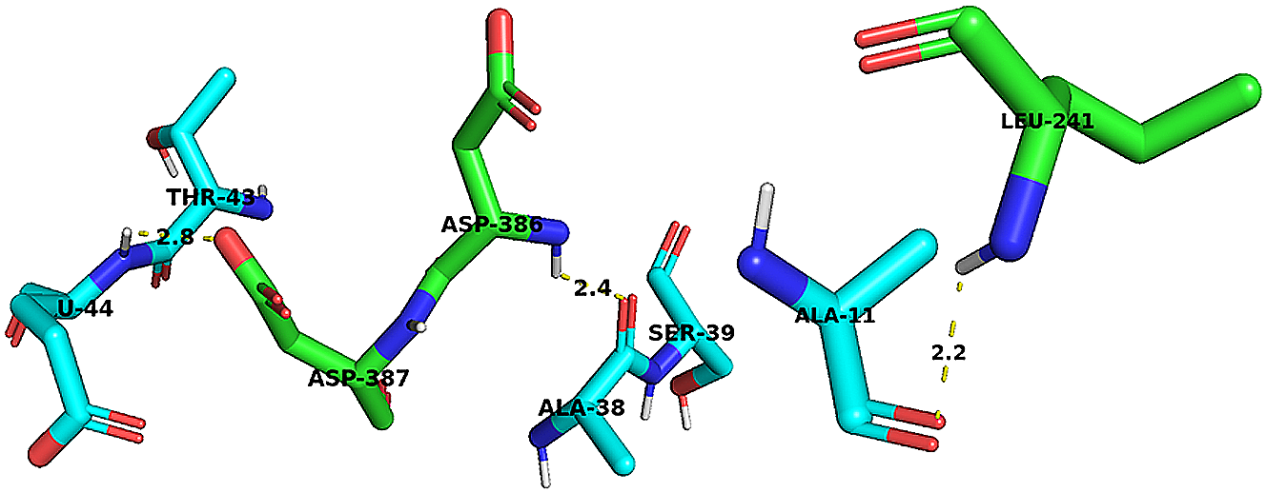 | | | | | | | | | | | | | | | | | | | | | | | | | | | | | | | | | |
| **Hydrogen bonds** | | | **Chain A** | | | | | | | | | | | | | | | **Chain B** | | | | | | | | | | | | | | | **Distance** |
|  |  |  | **S. No.** | | | | **Atom**  **number** | | | **Atom name** | | **Res. name** | | **Res. number** | | | | **Atom number** | | | | **Atom name** | | | **Res. name** | | | | **Res. number** | | | |  |
|  |  |  | 1. | | | | 1785 | | | O | | ARG | | 227 | | | | 5761 | | | | OG | | | SER | | | | 39 | | | | 2.54 |
|  |  |  | 2. | | | | 1890 | | | N | | LEU | | 241 | | | | 5558 | | | | O | | | ALA | | | | 11 | | | | 2.93 |
|  |  |  | 3. | | | | 1905 | | | OG | | SER | | 243 | | | | 5528 | | | | O | | | ILE | | | | 7 | | | | 3.09 |
|  |  |  | 4. | | | | 3003 | | | N | | ASP | | 386 | | | | 5758 | | | | O | | | ALA | | | | 38 | | | | 3.17 |
| **Salt bridge** | | | 1. | | | | 3029 | | | OD2 | | ASP | | 389 | | | | 5732 | | | | NZ | | | LYS | | | | 34 | | | | 3.86 |
| **Tau-Phosphoglyceromutase 78** | | | | | | | | | | | | | | | | | | | | | | | | | | | | | | | | | |
| 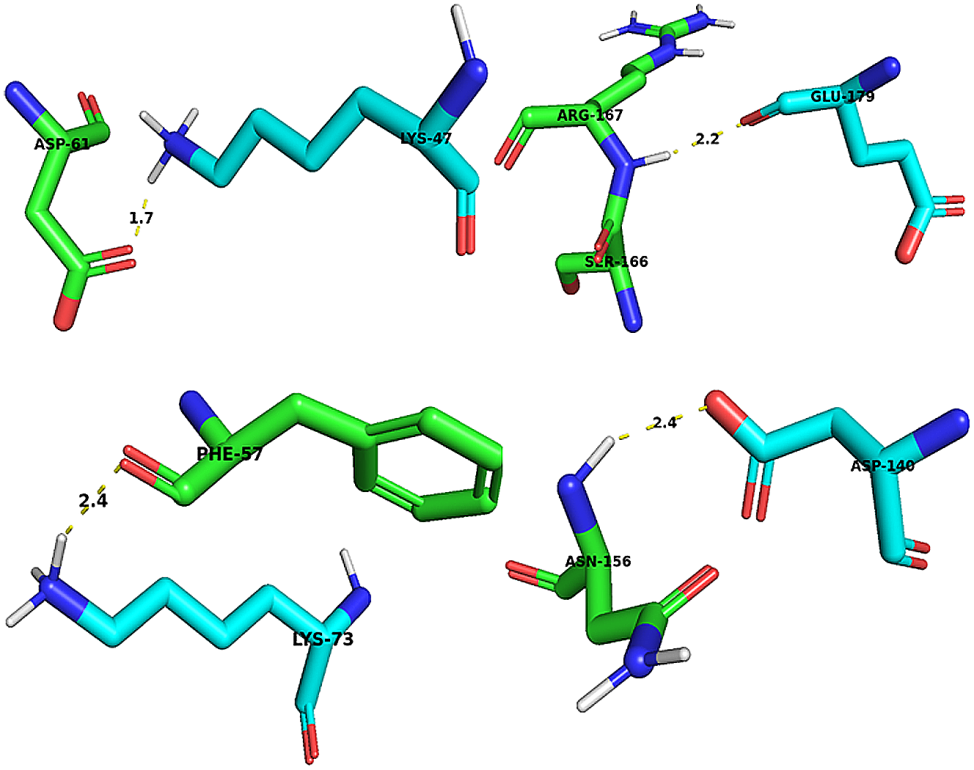 | | | | | | | | | | | | | | | | | | | | | | | | | | | | | | | | | |
| **Hydrogen bonds** | | | 1. | | | | 486 | | | OD1 | | ASP | | 61 | | | | 5850 | | | | NZ | | | LYS | | | | 47 | | | | 2.54 |
|  |  |  | 2. | | | | 1244 | | | N | | ASN | | 156 | | | | 6588 | | | | OD2 | | | ASP | | | | 140 | | | | 3.33 |
|  |  |  | 3. | | | | 1328 | | | N | | ARG | | 167 | | | | 6915 | | | | O | | | GLU | | | | 179 | | | | 3.19 |
|  |  |  | 4. | | | | 4134 | | | N | | GLY | | 539 | | | | 6318 | | | | O | | | TYR | | | | 108 | | | | 1.72 |
| **Salt bridges** | | | 1. | | | | 486 | | | OD1 | | ASP | | 61 | | | | 5850 | | | | NZ | | | LYS | | | | 47 | | | | 2.54 |
|  |  |  | 2. | | | | 1272 | | | NZ | | LYS | | 160 | | | | 6632 | | | | OE1 | | | GLU | | | | 145 | | | | 1.18 |
| **Tau-Alcohol dehydrogenase** | | | | | | | | | | | | | | | | | | | | | | | | | | | | | | | | | |
| 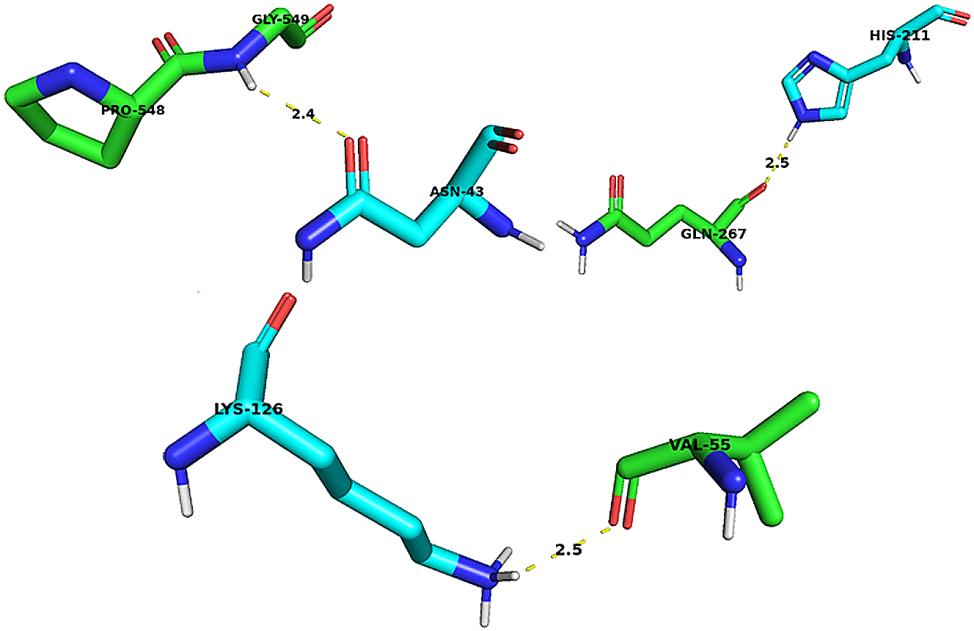 | | | | | | | | | | | | | | | | | | | | | | | | | | | | | | | | | |
| **Hydrogen bonds** | | | 1. | | | | | 437 | | O | | VAL | | 55 | | | | 6457 | | | | NZ | | | LYS | | | | 126 | | | | 2.99 |
|  |  |  | 2. | | | | | 2085 | | O | | GLN | | 267 | | | | 7089 | | | | NE2 | | | HIS | | | | 211 | | | | 3.17 |
|  |  |  | 3. | | | | | 4164 | | ND2 | | ASN | | 543 | | | | 5640 | | | | OD2 | | | ASP | | | | 22 | | | | 1.74 |
|  |  |  | 4. | | | | | 4164 | | ND2 | | ASN | | 543 | | | | 7121 | | | | OG | | | SER | | | | 216 | | | | 2.57 |
|  |  |  | 5. | | | | | 4179 | | OH | | TYR | | 546 | | | | 5815 | | | | ND2 | | | ASN | | | | 43 | | | | 1.78 |
|  |  |  | 6. | | | | | 4208 | | N | | GLY | | 549 | | | | 5809 | | | | OD1 | | | ASN | | | | 43 | | | | 3.22 |
| **Salt bridge** | | | 1. | | | | | 1282 | | OD2 | | ASP | | 161 | | | | 6082 | | | | NZ | | | LYS | | | | 78 | | | | 3.83 |
| **Tau-Mitochondrial Ribosomal protein L2** | | | | | | | | | | | | | | | | | | | | | | | | | | | | | | | | | |
| 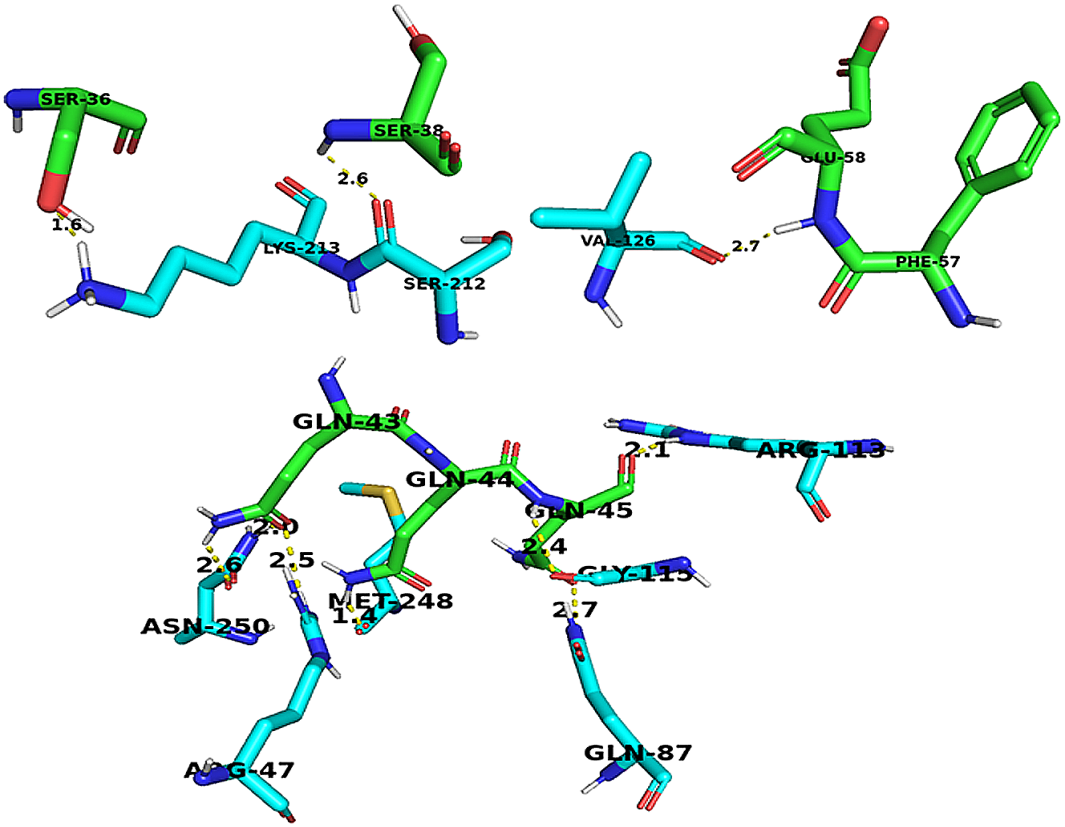 | | | | | | | | | | | | | | | | | | | | | | | | | | | | | | | | | |
| **Hydrogen bonds** | | | 1. | | | 112 | | | | | O | | SER | | 15 | | | | | 7468 | | | | OG | | SER | | | 254 | | | | 3.19 |
|  |  |  | 2. | | | 118 | | | | | O | | PRO | | 16 | | | | | 7476 | | | | N | | GLY | | | 255 | | | | 2.88 |
|  |  |  | 3. | | | 282 | | | | | OG | | SER | | 36 | | | | | 7139 | | | | NZ | | LYS | | | 213 | | | | 2.32 |
|  |  |  | 4. | | | 297 | | | | | N | | SER | | 38 | | | | | 7131 | | | | O | | SER | | | 212 | | | | 3.34 |
|  |  |  | 5. | | | 333 | | | | | OE1 | | GLN | | 43 | | | | | 7437 | | | | ND2 | | ASN | | | 250 | | | | 2.94 |
|  |  |  | 6. | | | 347 | | | | | NE2 | | GLN | | 44 | | | | | 7422 | | | | O | | MET | | | 248 | | | | 2.31 |
|  |  |  | 7. | | | 354 | | | | | N | | GLN | | 45 | | | | | 6390 | | | | O | | GLY | | | 115 | | | | 3.28 |
|  |  |  | 8. | | | 353 | | | | | O | | GLN | | 45 | | | | | 6373 | | | | NE | | ARG | | | 113 | | | | 3.05 |
|  |  |  | 9. | | | 453 | | | | | O | | PHE | | 57 | | | | | 6486 | | | | N | | GLU | | | 129 | | | | 2.35 |
|  |  |  | 10. | | | 487 | | | | | OD2 | | ASP | | 61 | | | | | 6273 | | | | OG1 | | THR | | | 100 | | | | 2.08 |
|  |  |  | 11. | | | 489 | | | | | OG | | SER | | 62 | | | | | 6644 | | | | NH1 | | ARG | | | 149 | | | | 1.69 |
| **Salt Bridges** | | | 1. | | | 542 | | | | | NZ | | LYS | | 68 | | | | | 7720 | | | | OE2 | | GLU | | | 285 | | | | 3.96 |
|  |  |  | 2. | | | 4993 | | | | | OE1 | | GLU | | 652 | | | | | 7711 | | | | NZ | | LYS | | | 284 | | | | 3.91 |
| **Tau-Retinin** | | | | | | | | | | | | | | | | | | | | | | | | | | | | | | | | | |
| 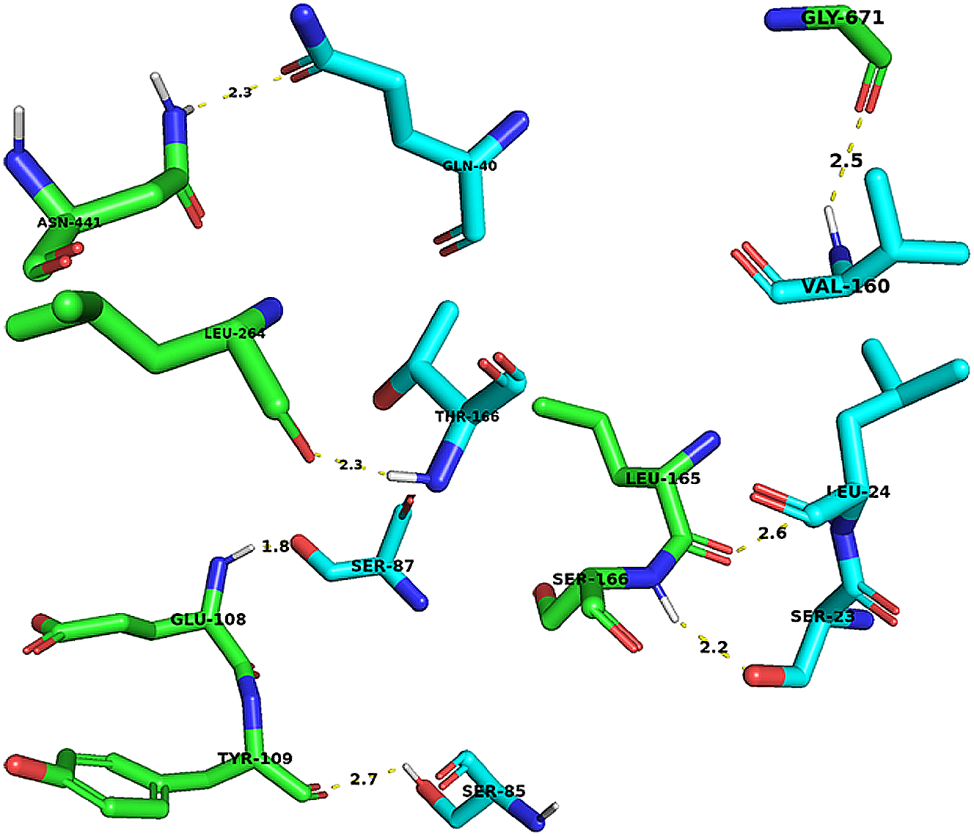 | | | | | | | | | | | | | | | | | | | | | | | | | | | | | | | | | |
| **Hydrogen bonds** | | 1. | | | 1312 | | | | | O | | LEU | | | | 165 | | 5644 | | | | | OG | | | SER | | 23 | | | | 3.23 | |
|  |  | 2. | | | 1312 | | | | | O | | LEU | | | | 165 | | 5654 | | | | | N | | | LEU | | 24 | | | | 3.18 | |
|  |  | 3. | | | 1321 | | | | | N | | SER | | | | 166 | | 5644 | | | | | OG | | | SER | | 23 | | | | 3.19 | |
|  |  | 4. | | | 1388 | | | | | OG | | SER | | | | 175 | | 5744 | | | | | O | | | VAL | | 36 | | | | 2.73 | |
|  |  | 5. | | | 2061 | | | | | O | | LEU | | | | 264 | | 6685 | | | | | N | | | THR | | 166 | | | | 3.33 | |
|  |  | 6. | | | 2077 | | | | | O | | MET | | | | 266 | | 6678 | | | | | NH2 | | | ARG | | 165 | | | | 2.72 | |
|  |  | 7. | | | 3414 | | | | | ND2 | | ASN | | | | 441 | | 5771 | | | | | OE1 | | | GLN | | 40 | | | | 2.94 | |
|  |  | 8. | | | 3447 | | | | | OD2 | | ASP | | | | 446 | | 5815 | | | | | OG | | | SER | | 46 | | | | 2.27 | |
| **Salt Bridge** | | 1. | | | 4434 | | | | | NZ | | LYS | | | | 578 | | 6114 | | | | | OD1 | | | ASP | | 89 | | | | 3.27 | |
| **Tau-Globin** | | | | | | | | | | | | | | | | | | | | | | | | | | | | | | | | | |
| 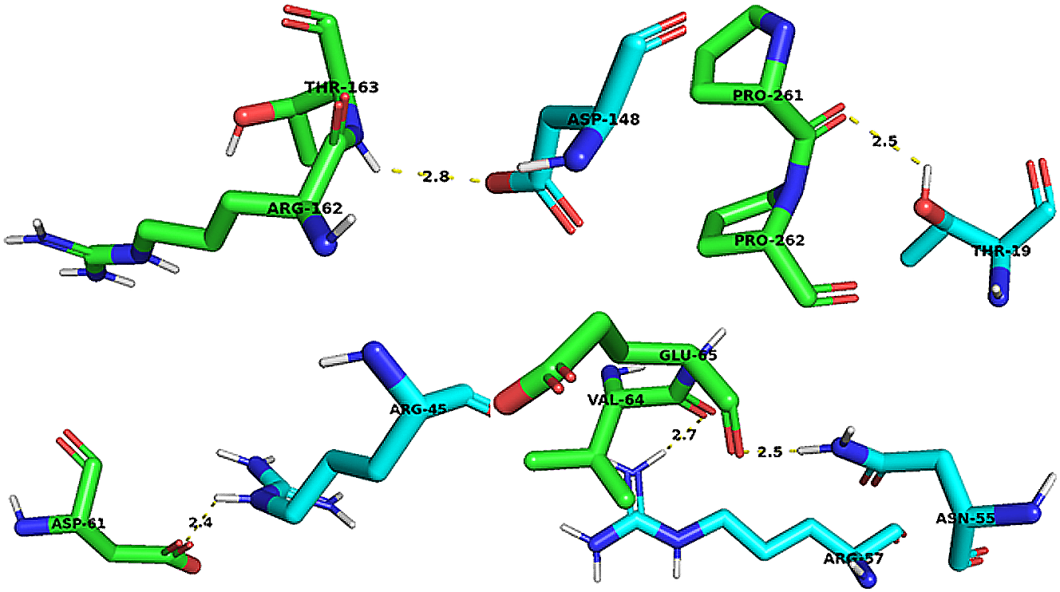 | | | | | | | | | | | | | | | | | | | | | | | | | | | | | | | | | |
| **Hydrogen bonds** | 1. | | | 486 | | | | | OD1 | | | | ASP | | | | 61 | | 5842 | | NE | | | | | | ARG | | | 45 | 2.89 | | |
|  | 2. | | | 486 | | | | | OD1 | | | | ASP | | | | 61 | | 5847 | | NH2 | | | | | | ARG | | | 45 | 2.82 | | |
|  | 3. | | | 504 | | | | | N | | | | VAL | | | | 64 | | 5824 | | O | | | | | | PRO | | | 43 | 2.58 | | |
|  | 4. | | | 511 | | | | | O | | | | GLU | | | | 65 | | 5922 | | ND2 | | | | | | ASN | | | 55 | 2.90 | | |
|  | 5. | | | 2038 | | | | | O | | | | PRO | | | | 261 | | 5632 | | OG1 | | | | | | THR | | | 19 | 3.08 | | |
|  | 6. | | | 2045 | | | | | O | | | | PRO | | | | 262 | | 5632 | | OG1 | | | | | | THR | | | 19 | 2.74 | | |
|  | 7. | | | 2052 | | | | | O | | | | GLN | | | | 263 | | 5628 | | N | | | | | | ALA | | | 18 | 3.16 | | |
| **Salt Bridges** | 1. | | | 486 | | | | | OD1 | | | | ASP | | | | 61 | | 5847 | | NH2 | | | | | | ARG | | | 45 | 2.82 | | |
|  | 2. | | | 1281 | | | | | OD1 | | | | ASP | | | | 161 | | 6687 | | N2 | | | | | | LYS | | | 153 | 3.73 | | |

**Table S2:** Pictorial Representation of the of the hydrogen bonds interactions both as the differential colour surfaces as overall in the section A (Chain B) with Tau Protein (Chain A) and identification of Tau pocket to host the identified proteins (as depicted in the section B) of *Drosophila melanogaster.*

| **Interaction type** | **Interchain Hydrogen Bonding Interaction Surface** |
| --- | --- |
| **Tau-OBP44a** | 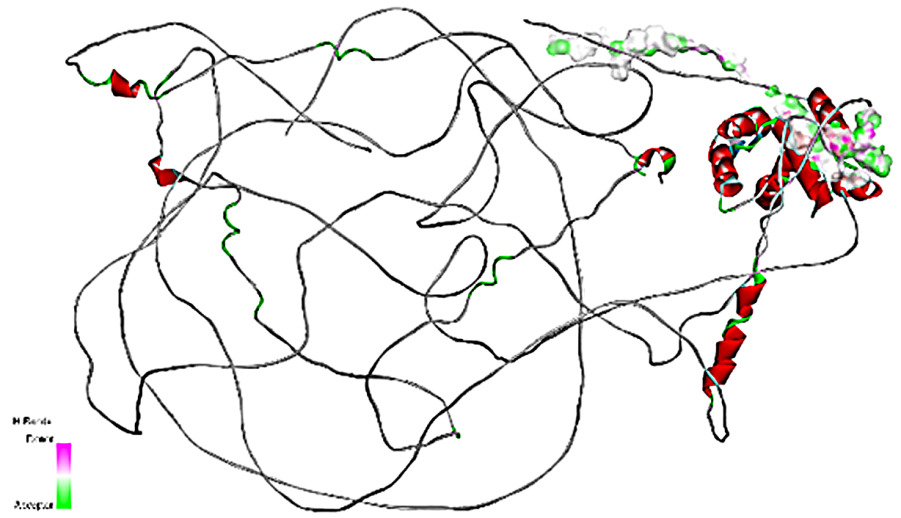  **(A)** |
|  | 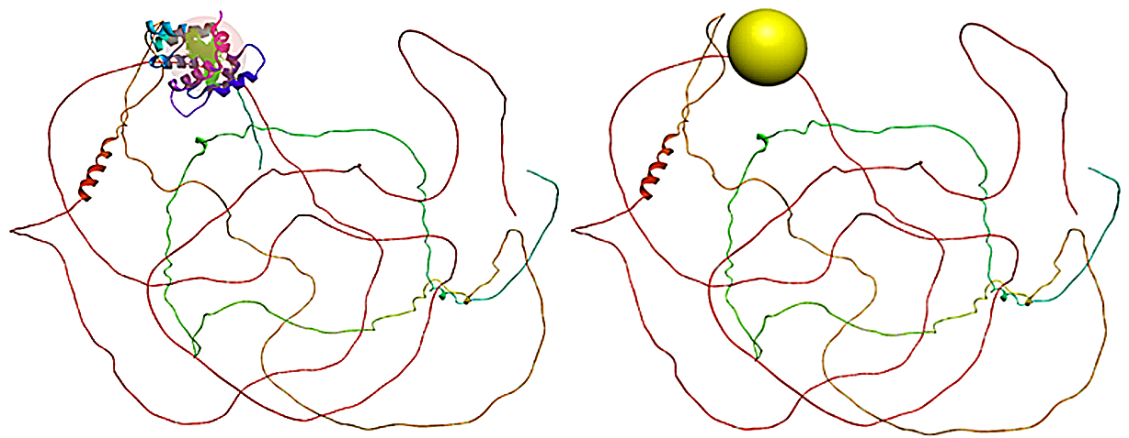 |
| **Tau-Phosphogly-ceromutase 78** | 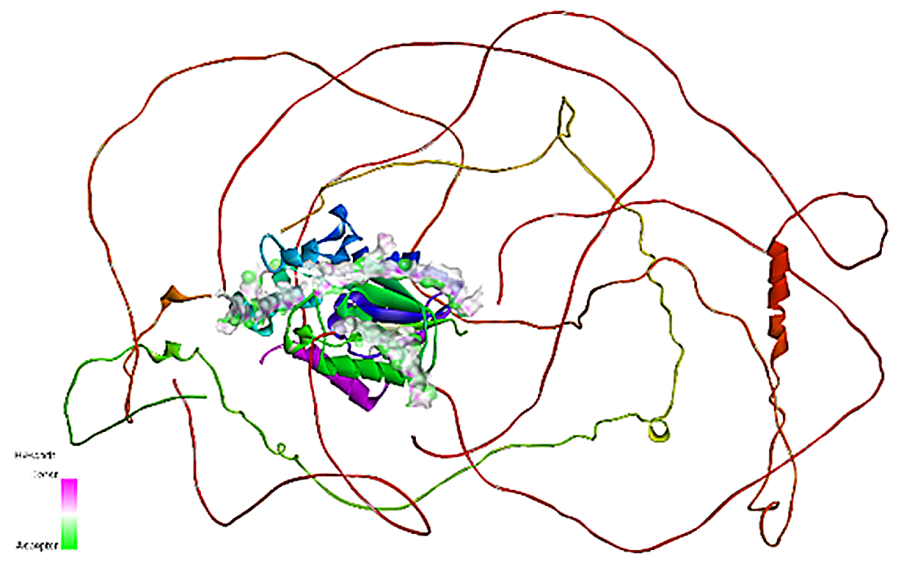  **(A)** |
|  | 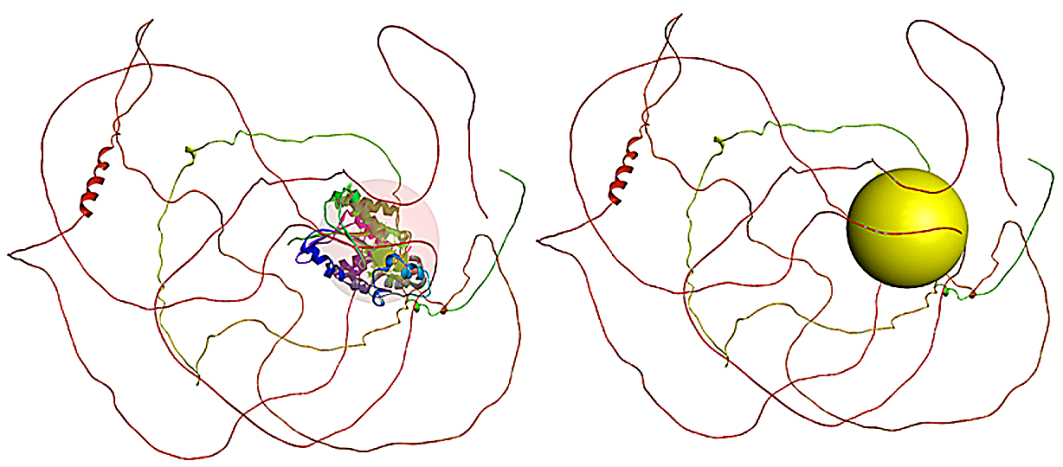  **(B)** |
| **Tau-Alcohol Dehydrogenase** | 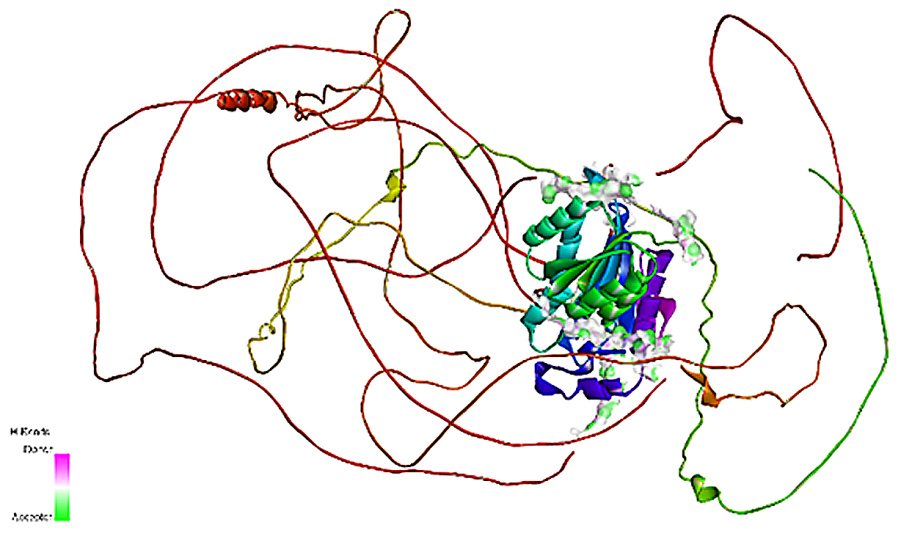  **(A)** |
|  | 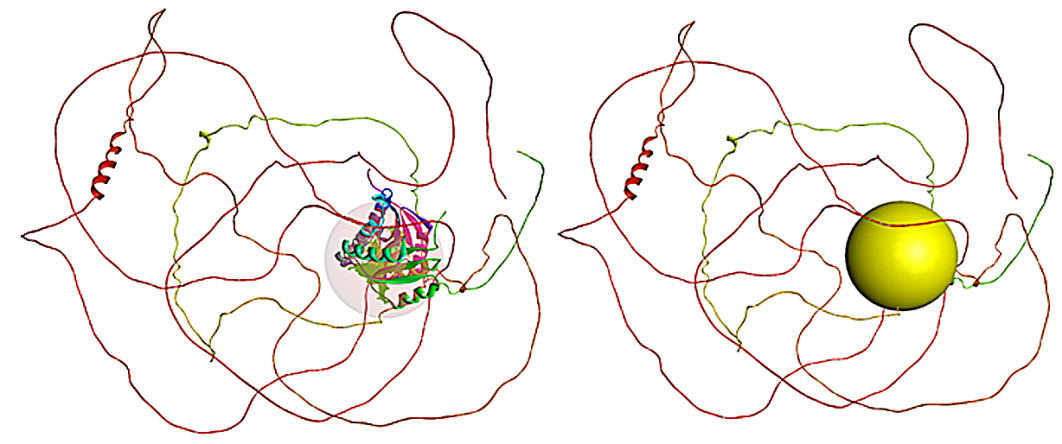  **(B)** |
| **Tau-Mitochondrial Ribosomal protein L2** | 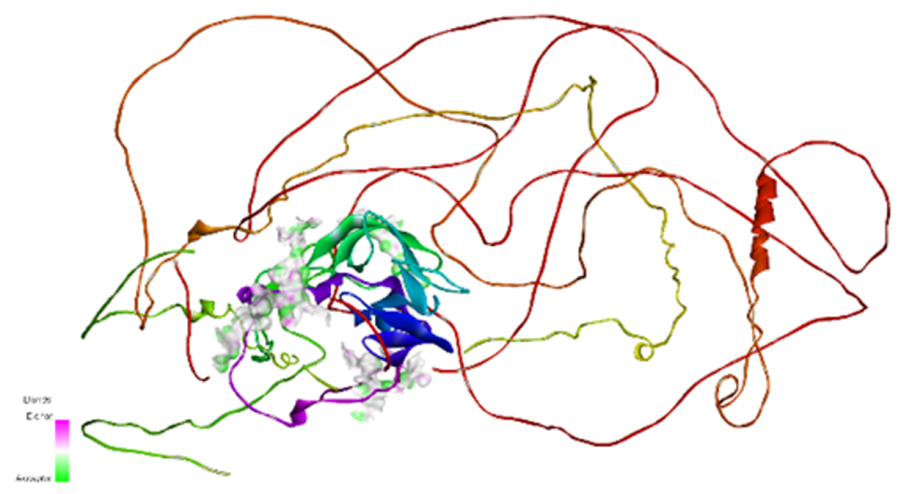  **(A)** |
|  | 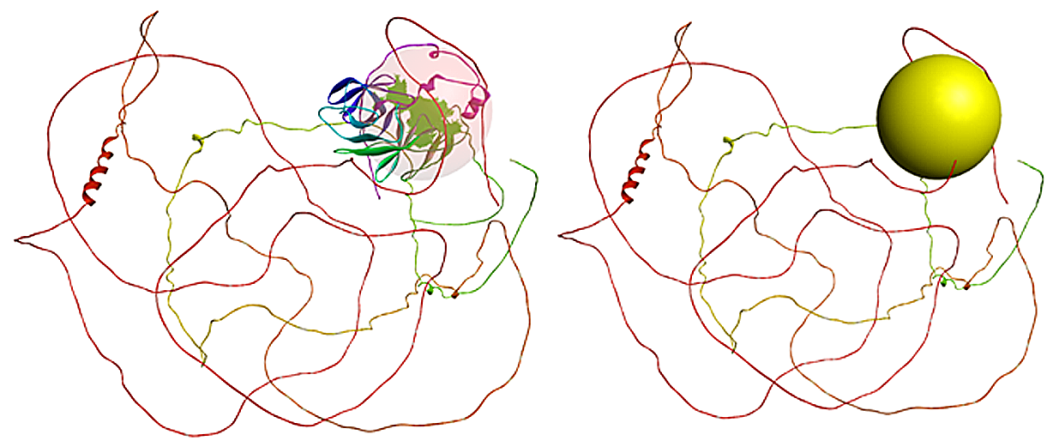  **(B)** |
| **Tau-Retinin** | 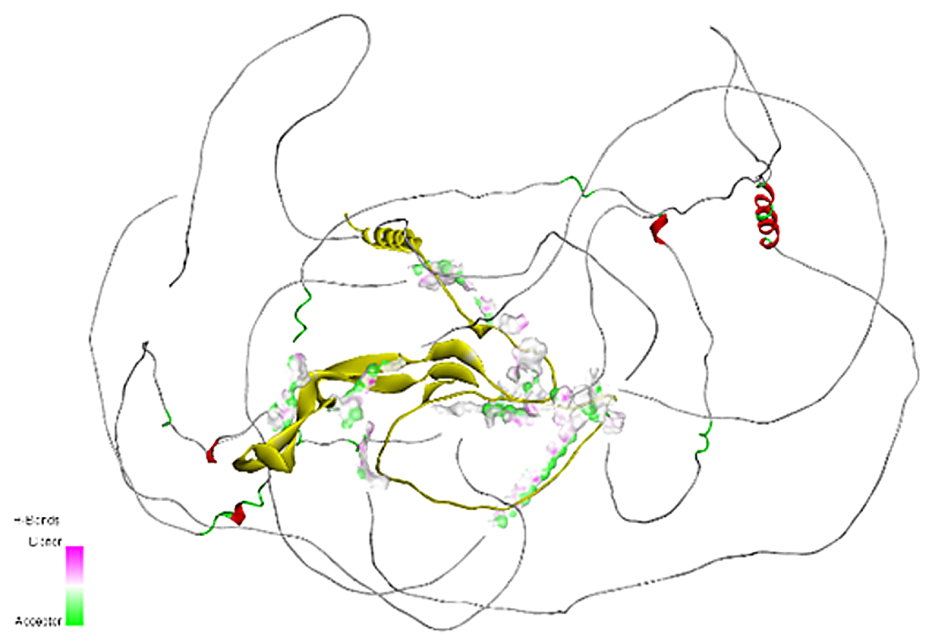  **(A)** |
|  | 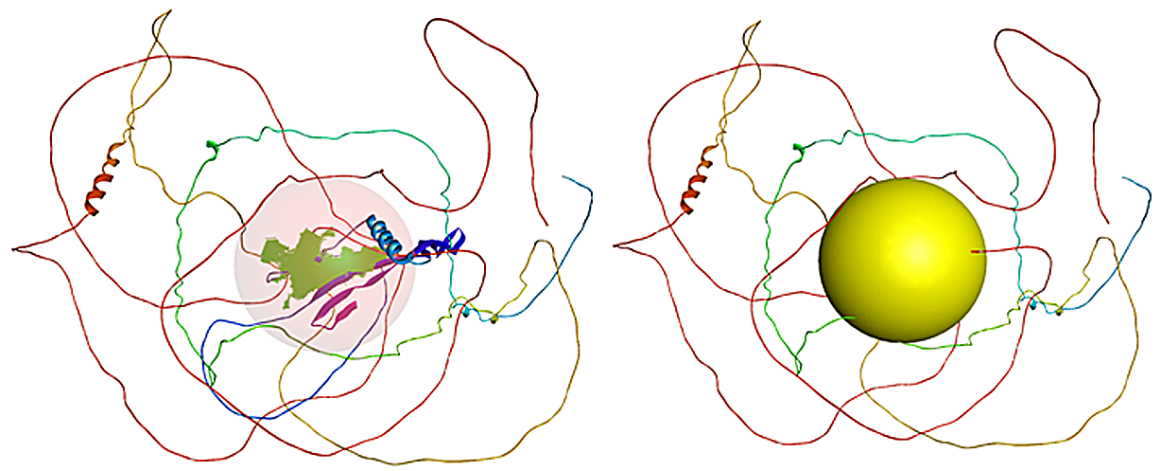  **(B)** |
| **Tau-Globin** | 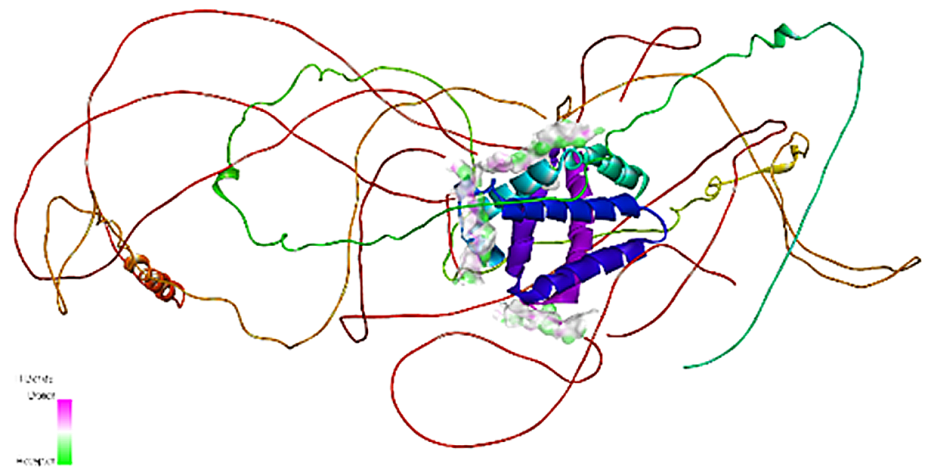  **(A)** |
|  | 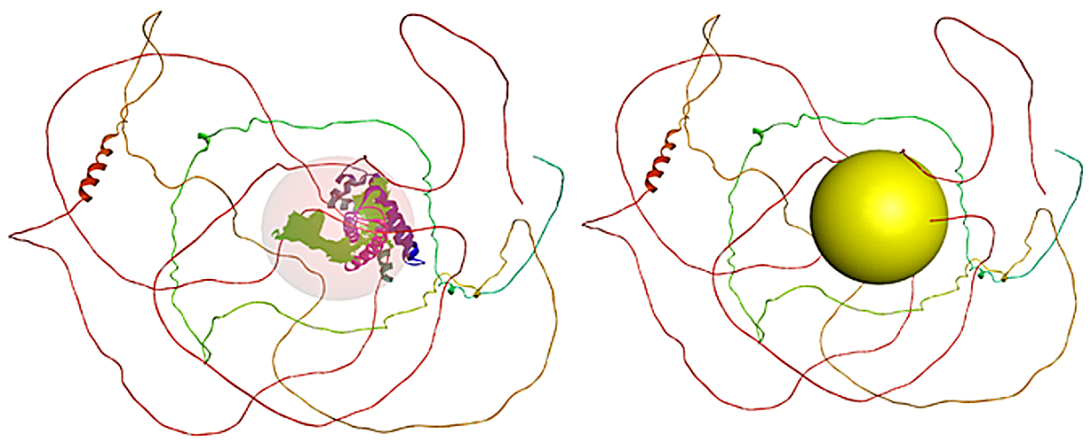  **(B)** |

**Table S3:** Pictorial Representation of the all-different type of interactions of identified proteins (as depicted in the last column) (Chain B) with Tau Protein (Chain A) of *Drosophila melanogaster* as depicted in the middle column*.* The corresponding Ramachandran Plots for the interactions for the same is depicted in last column with their corresponding statistics.

| **Interaction type** | **Interchain Interactions of Chain A and B** | **Corresponding Ramachandran plots for the interactions** |
| --- | --- | --- |
| **Tau-OBP44a** | 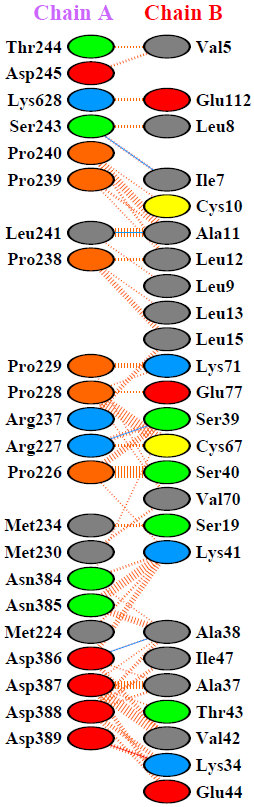  \| **Key:** \| 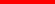 \| \| Salt bridges \|; \| \| 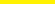 \| \| Disulphide bonds \|; \| \| 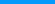 \| \| Hydrogen bonds \|; \| \| 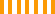 \| \| Non-bonded contacts \| | 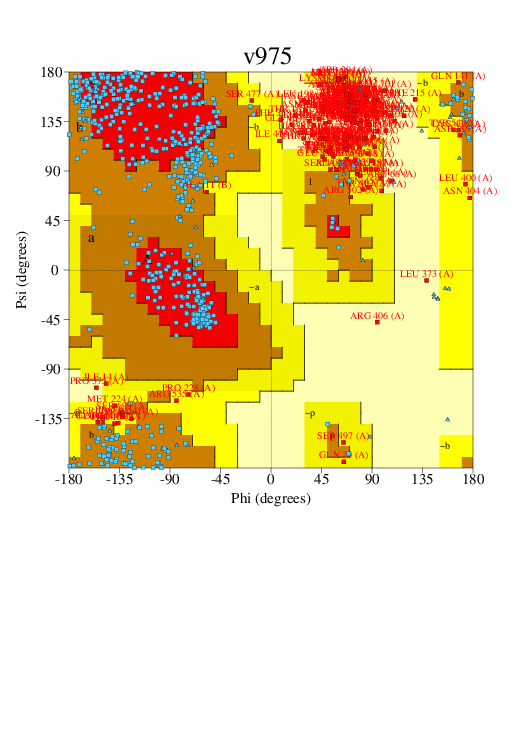 **Ramachandran Plot statistics** Attributes \| No. of residues \| %; Most favoured regions [A,B,L] \| 330 \| 48.1; Additional allowed regions [a,b,l,p] \| 179 \| 26.1; Generously allowed regions [~a,~b,~l,~p] \| 66 \| 9.6; Disallowed regions [XX] \| 111 \| 16.2; Non-glycine and non-proline residues \| 686 \| 100; End-residues (excl. Gly and Pro) \| 4 \|; Glycine residues \| 56 \|; Proline residues \| 114 \|; Total number of residues \| 860 \| |
| **Tau-Phosphogly-ceromutase 78** | 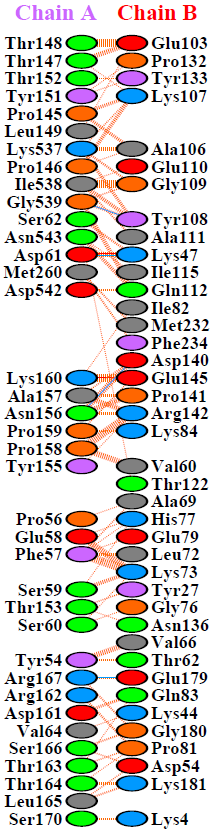  \| **Key:** \| 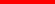 \| \| Salt bridges \|; \| \| 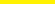 \| \| Disulphide bonds \|; \| \| 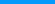 \| \| Hydrogen bonds \|; \| \| 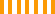 \| \| Non-bonded contacts \| | 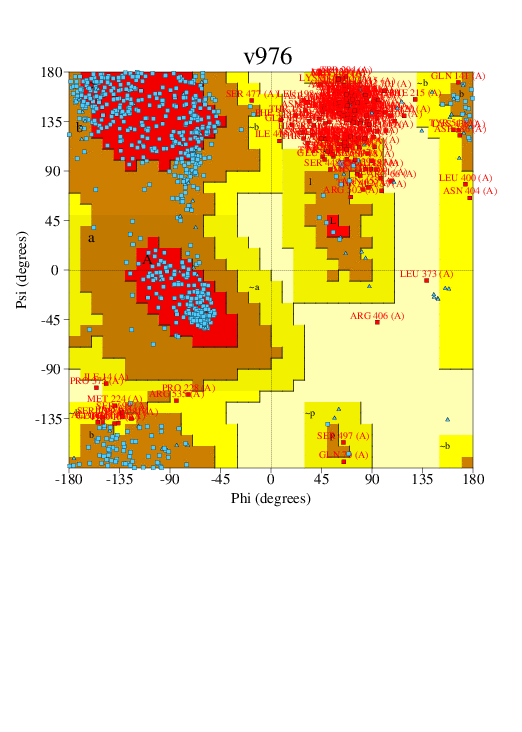 **Ramachandran Plot statistics** Attributes \| No. of residues \| %; Most favoured regions [A,B,L] \| 419 \| 54.3; Additional allowed regions [a,b,l,p] \| 178 \| 23.1; Generously allowed regions [~a,~b,~l,~p] \| 63 \| 8.2; Disallowed regions [XX] \| 111 \| 14.4; Non-glycine and non-proline residues \| 771 \| 100; End-residues (excl. Gly and Pro) \| 4 \|; Glycine residues \| 70 \|; Proline residues \| 127 \|; Total number of residues \| 972 \| |
| **Tau-Alcohol Dehydrogenase** | 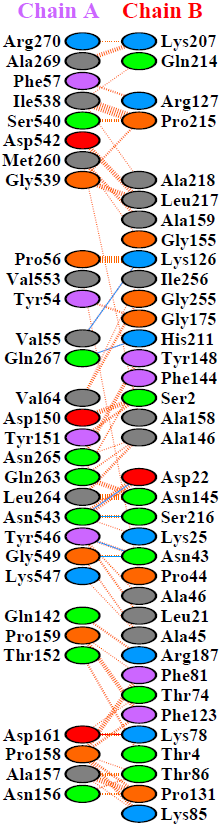  \| **Key:** \| 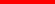 \| \| Salt bridges \|; \| \| 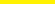 \| \| Disulphide bonds \|; \| \| 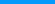 \| \| Hydrogen bonds \|; \| \| 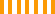 \| \| Non-bonded contacts \| | 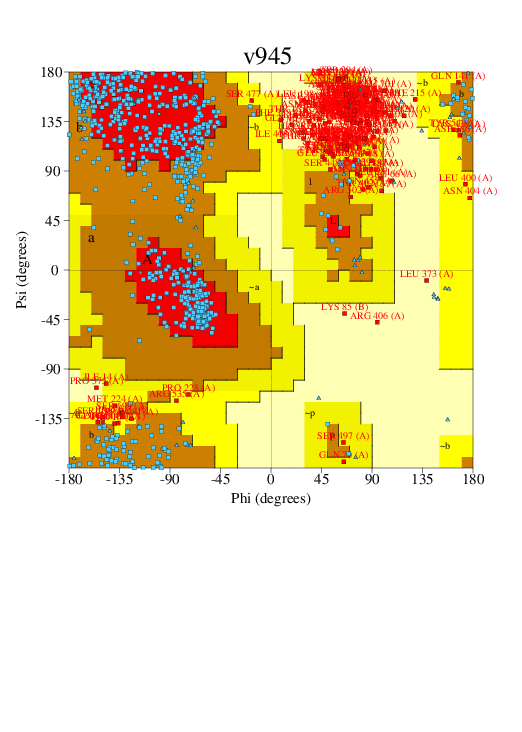 **Ramachandran Plot statistics** Attributes \| No. of residues \| %; Most favoured regions [A,B,L] \| 422 \| 54.2; Additional allowed regions [a,b,l,p] \| 181 \| 23.3; Generously allowed regions [~a,~b,~l,~p] \| 63 \| 8.1; Disallowed regions [XX] \| 112 \| 14.4; Non-glycine and non-proline residues \| 778 \| 100; End-residues (excl. Gly and Pro) \| 4 \|; Glycine residues \| 69 \|; Proline residues \| 122 \|; Total number of residues \| 973 \| |
| **Tau-Mitochondrial Ribosomal protein L2** | 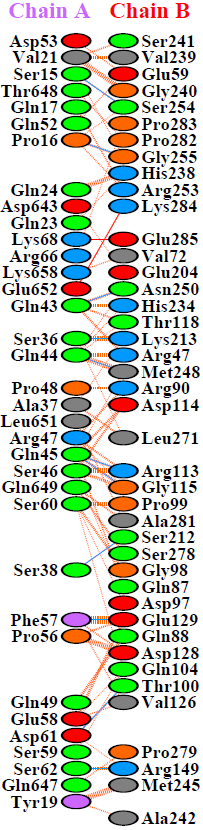  \| **Key:** \| 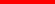 \| \| Salt bridges \|; \| \| 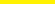 \| \| Disulphide bonds \|; \| \| 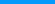 \| \| Hydrogen bonds \|; \| \| 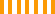 \| \| Non-bonded contacts \| | 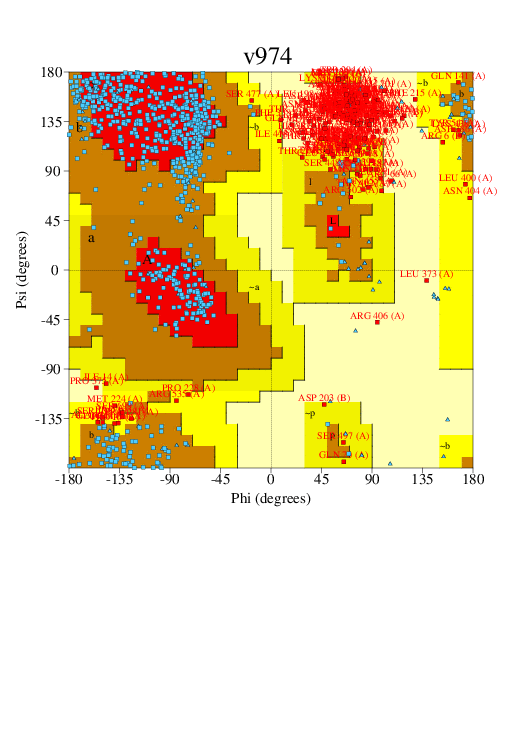 **Ramachandran Plot statistics** Attributes \| No. of residues \| %; Most favoured regions [A,B,L] \| 422 \| 53.0; Additional allowed regions [a,b,l,p] \| 189 \| 23.7; Generously allowed regions [~a,~b,~l,~p] \| 69 \| 8.7; Disallowed regions [XX] \| 116 \| 14.6; Non-glycine and non-proline residues \| 796 \| 100; End-residues (excl. Gly and Pro) \| 4 \|; Glycine residues \| 76 \|; Proline residues \| 135 \|; Total number of residues \| 1011 \| |
| **Tau-Retinin** | 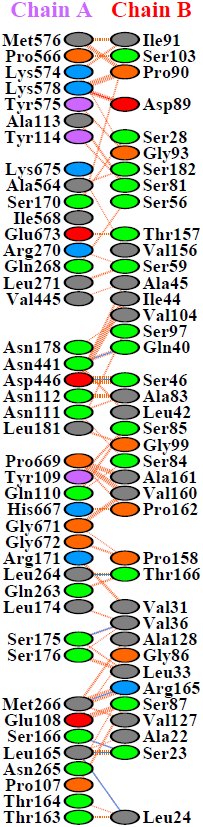  \| **Key:** \| 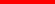 \| \| Salt bridges \|; \| \| 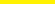 \| \| Disulphide bonds \|; \| \| 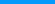 \| \| Hydrogen bonds \|; \| \| 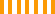 \| \| Non-bonded contacts \| | 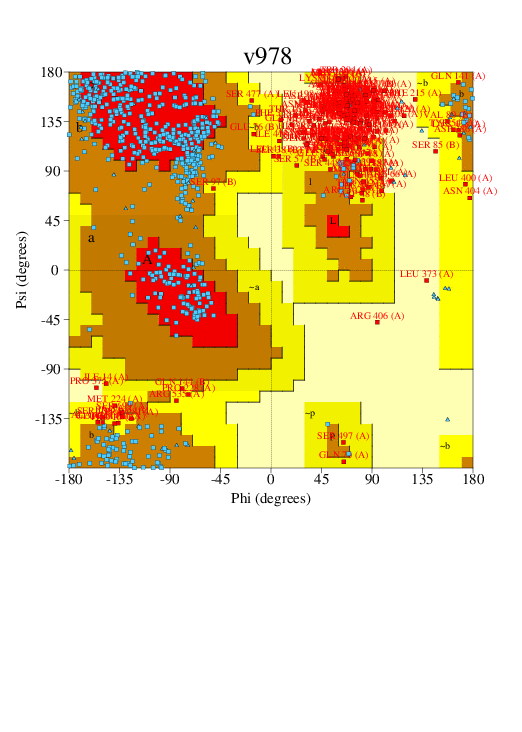 **Ramachandran Plot statistics** Attributes \| No. of residues \| %; Most favoured regions [A,B,L] \| 343 \| 47.6; Additional allowed regions [a,b,l,p] \| 189 \| 26.2; Generously allowed regions [~a,~b,~l,~p] \| 74 \| 10.3; Disallowed regions [XX] \| 114 \| 15.8; Non-glycine and non-proline residues \| 720 \| 100; End-residues (excl. Gly and Pro) \| 4 \|; Glycine residues \| 58 \|; Proline residues \| 126 \|; Total number of residues \| 908 \| |
| **Tau-Globin** | 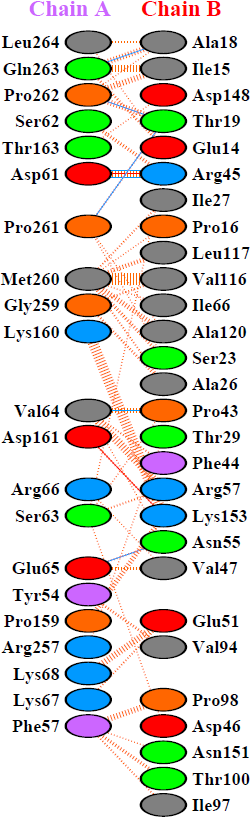  \| **Key:** \| 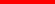 \| \| Salt bridges \|; \| \| 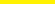 \| \| Disulphide bonds \|; \| \| 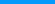 \| \| Hydrogen bonds \|; \| \| 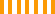 \| \| Non-bonded contacts \| | 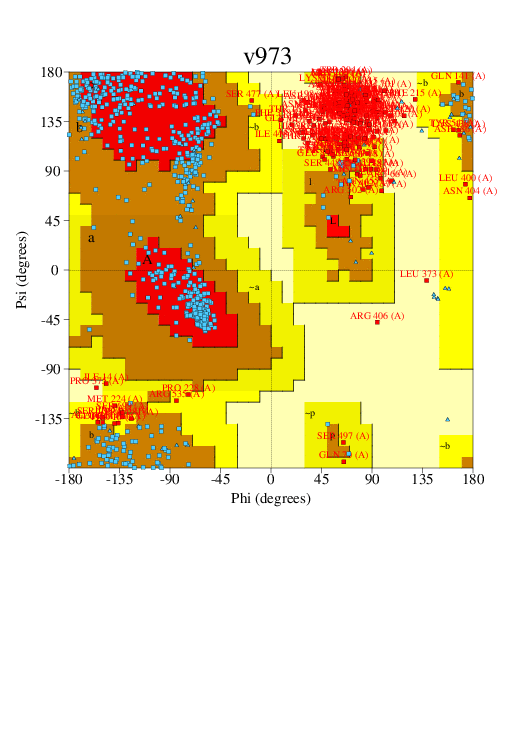 **Ramachandran Plot statistics** Attributes \| No. of residues \| %; Most favoured regions [A,B,L] \| 352 \| 50.9; Additional allowed regions [a,b,l,p] \| 165 \| 23.9; Generously allowed regions [~a,~b,~l,~p] \| 63 \| 9.1; Disallowed regions [XX] \| 111 \| 16.1; Non-glycine and non-proline residues \| 691 \| 100; End-residues (excl. Gly and Pro) \| 4 \|; Glycine residues \| 58 \|; Proline residues \| 117 \|; Total number of residues \| 870 \| |
